# Virus-mediated tau aggregate seeding in a cellular model

**DOI:** 10.64898/2026.06.25.734433

**Authors:** Patricia Yuste-Checa, Roman D. Martinez-Piera, Freya Herrmann-Sim, L. Sofia Ronquillo-Silva, Roman Körner, F. Ulrich Hartl

## Abstract

Formation of neuronal tau protein aggregates is a defining feature of tauopathies, including Alzheimer’s disease and frontotemporal dementia. Tau pathology propagates across brain regions by a cell-to-cell aggregate seeding mechanism. While epidemiological and experimental studies over the past three decades have implicated viral infections in aggregate-associated neurodegeneration, the underlying mechanisms remain unclear. Here, we show in a cell culture model that seeding competent tau aggregates are efficiently packaged into lentiviral particles and induce tau aggregation in recipient cells in a virus receptor-dependent manner. Tau aggregates interact with the viral Gag polyprotein, likely facilitating their incorporation into virions. Virus-dependent tau seeding requires protease-mediated maturation of the envelope protein to enable efficient fusion of the viral envelope with the plasma membrane of the recipient cell. Thus, tau aggregate packaging is compatible with the formation of mature, infectious virions. These findings are consistent with a possible role of viral infection in neurodegenerative disease progression and offer a robust, versatile platform to model tau propagation in cellular systems and animal models.

## Introduction

The major neurodegenerative diseases (NDs), including Alzheimer’s (AD) and Parkinson’s disease, are classified as proteinopathies, defined by the accumulation of specific disease proteins in intra- or extracellular aggregates with a characteristic fibrillar structure referred to as amyloid^1–3^. Aggregates of the microtubule binding protein tau accumulate in various tauopathies, including AD. The amyloid fibrils associated with NDs are considered “prion-like”, owing to their ability to “seed” aggregation of their native, monomeric counterparts, resulting in the self-propagation of pathological aggregates^1,4^. Seed aggregates can travel from cell to cell through several defined mechanisms, including direct transfer via tunnelling nanotubes, release into the extracellular space or secretion within extracellular vesicles followed by uptake by recipient cells. Once internalized by naïve cells, these aggregates may template the aggregation of the monomeric protein, a process thought to underlie disease progression^1,5,6^.

Most cases of NDs are sporadic, with approximately 85-95% occurring in the absence of a clearly identifiable single genetic cause. These disorders are therefore thought to be the result of a cumulative interplay of multiple factors, including incompletely penetrant risk alleles, aging-related cellular proteostasis decline, and environmental risk factors^7–9^. One notable environmental risk factor is viral infection. The first evidence linking viral infection to neurodegeneration emerged 35 years ago, with the detection of herpes simplex virus type 1 (HSV-1) DNA in the brains of patients with AD^10,11^. Since then, a growing body of evidence has implicated viral infection as a potential contributor to sporadic forms of neurodegeneration, supported by epidemiological associations, experimental data and the observation that vaccination is associated with reduced disease risk^12–20^.

Viruses may contribute to neurodegeneration through multiple, mutually non-exclusive mechanisms, such as induction of chronic neuroinflammation, enhanced expression of amyloidogenic proteins, promotion of amyloid aggregation, dysregulation of the proteostasis network^18,21^ or transfer of seed aggregates by infection. Consistent with the latter possibility, exosome vesicles have been shown to contain aggregate seeds^22^ and vesicles produced after infection^23^ or decorated with viral envelope proteins can facilitate aggregate spread in cell culture^24,25^. Notably, extracellular vesicles share striking similarities with enveloped viruses. Both are surrounded by a lipid bilayer and can carry nucleic acid and protein cargoes to be ultimately delivered into recipient cells following fusion with the plasma membrane^26,27^. Additionally, the incorporation of host cellular proteins into some viruses is a well-established phenomenon relying either on non-specific passive incorporation or specific interactions between viral and host protein^28,29^. Despite the mounting epidemiological and experimental evidence associating viral infection and neurodegeneration, the causal significance of these findings remains unclear.

Here we show the efficient incorporation of pathological, seeding competent tau aggregates into lentiviral particles (LVs) in a cellular system and their ability to efficiently induce tau aggregation in infected cells in a virus receptor-dependent manner. To elucidate the underlying mechanisms of this virus-mediated tau aggregate seeding, we performed a detailed biochemical characterization of the viral particles. Our analysis revealed that the viral protease (PR) and the Gag polyprotein exhibit affinity for tau aggregates and may facilitate their incorporation into viral particles. We found that while the viral PR is dispensable for aggregate incorporation, it is essential for the LVs to acquire fusion competence with the cellular plasma membrane, primarily through proteolytic processing of the ecotropic Murine Leukemia Virus envelope (E-MLV Env, murine-tropic) protein used here to pseudotype the LVs. Therefore, fusion with the cellular plasma membrane achieved by viral PR activity is required for tau seeding in this system and importantly, packaging of tau seed aggregates is compatible with the formation of mature, infectious virions.

Collectively, our data support and extend previous evidence suggesting that viruses may contribute to the progression and pathogenesis of NDs, particularly by facilitating tau aggregate spreading. Moreover, the experimental system we describe represents a powerful and versatile platform to model tau aggregate-related neurodegeneration in human cellular models with high sensitivity and specificity.

## Results

### Lentiviruses produced in cells containing tau aggregates are seeding competent

To test whether pathological, seeding competent tau aggregates may be incorporated into viral particles, we produced lentiviruses (LVs, human immunodeficiency virus type 1, HIV-1 derived) in HEK293T cells containing aggregated or soluble tauRD (tau residues 244-371, P301L/V331M)-YFP (Fig. 1a). The aggregate-containing tauRD-YFP cell line was generated using tauRD-YFP aggregate seeds from the previously described Clone 9 cells^30^ that stably propagate tau amyloid. Cells were transfected with plasmids encoding for the structural viral proteins Gag/GagPol and mRuby with the lentiviral mRNA packing signal to monitor infection. Due to safety concerns about producing potentially infectious aggregate-loaded viruses, we pseudotyped the LVs to selectively infect murine cells by using the plasmid encoding for the ecotropic envelope protein from the Murine Leukemia Virus (E-MLV Env, murine-tropic). To monitor aggregate seeding competence of the LV particles, we used a reporter HEK293T cell line that co-expresses the FRET pair tauRD-YFP and tauRD-mTurquoise (tau residues 244-371 P301L/V331M; tau-YT)^31^. We murinized this cell line by overexpressing the murine receptor mCat1 (Tau-YTmCat1) (Fig. 1a and Supplementary Fig. 1a), which serves as the receptor for E-MLV Env used to pseudotype the LVs. LVs produced in cells containing soluble tau (LVTauS) were infectious (mRuby signal) but unable to induce aggregation (FRET signal) in the reporter cell line (Fig. 1b). Strikingly, viral particles produced in cells containing tau aggregates (LVTauAgg) efficiently “infected” the murinized reporter cells with aggregate pathology in a concentration dependent manner, suggesting that viral particles may be able to entrap seeding-competent tau aggregates (Fig. 1b). LVTauAgg particles were unable to infect or induce aggregation in the original reporter cell line (Tau-YT) which does not express the murine receptor mCat1 (Fig. 1c). Therefore, aggregation induction in the receptor cells by the viral particles produced in cells containing tau aggregates is dependent on the expression of the viral receptor.

**Figure 1.**
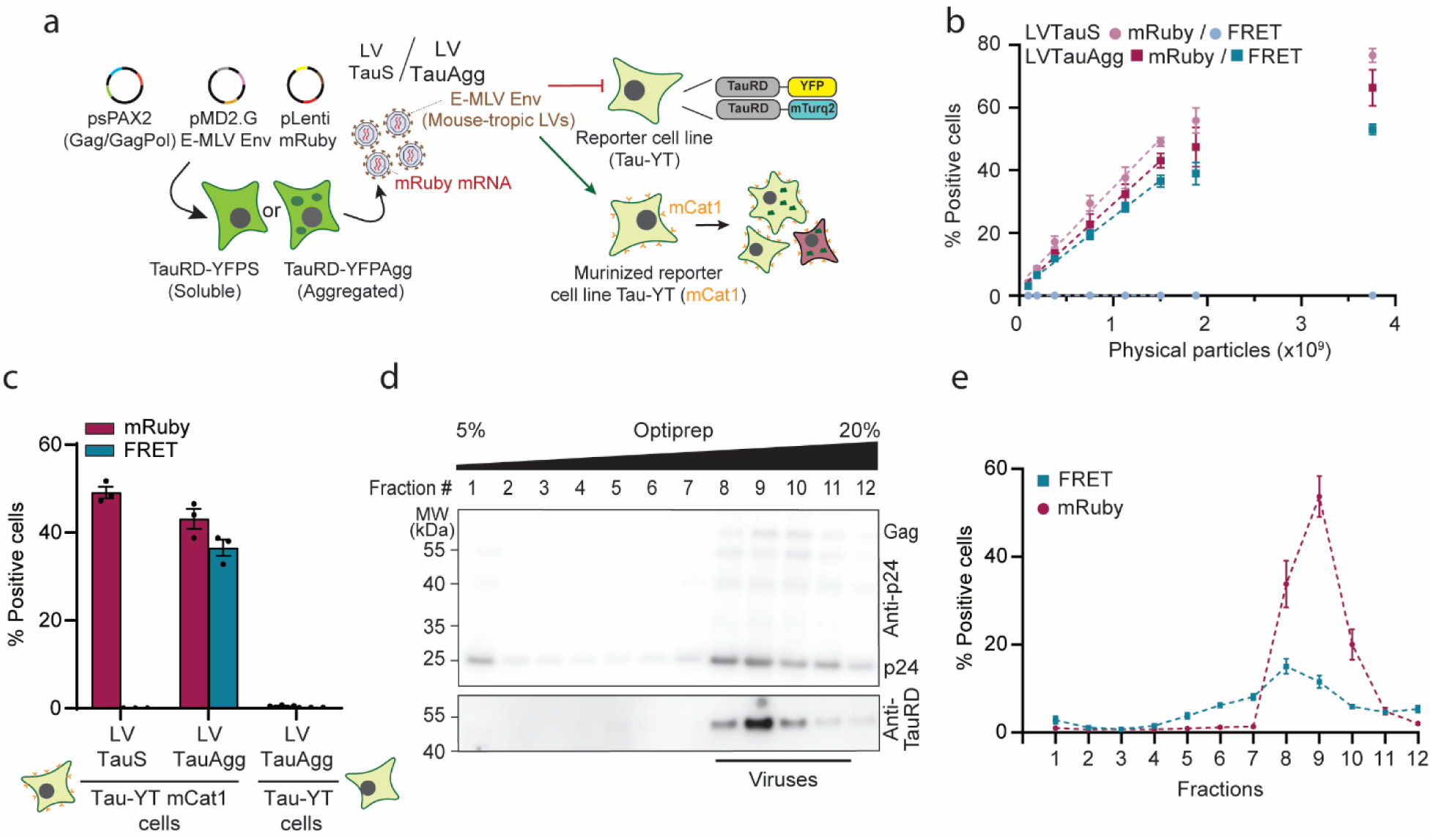
Viral particles produced in cells containing tau aggregates are seeding competent. **a.** Workflow of lentiviral production and seeding experiment. HEK293T cells expressing soluble (S) or aggregated (Agg) TauRD-YFP (P301L/V337M) were transfected with plasmids encoding for the structural viral proteins Gag/GagPol, the ecotropic Murine Leukemia Virus envelope protein (E-MLV Env, murine-tropic), and mRuby with the lentiviral mRNA packing signal. Lentiviruses (LVs) produced in cells with soluble or aggregated tau (LVTauS and LVTauAgg) were added to reporter cells co-expressing TauRD fused to the FRET pair of YFP and mTurquoise2 (TauRD-YT) and murinized by expression of the murine receptor mCat1 (Tau-YTmCat1), receptor for the E-MLV Env protein. **b.** Increasing amounts of the LVTauS and LVTauAgg were added to the murinized reporter cells (Tau-YTmCat1) and induction of aggregation (FRET positive cells, cyan tones) and infection (mRuby positive cells, maroon tones) was quantified by flow cytometry. Dashed lines represent linear fits to the data. Mean ± SEM (n = 3 biological replicates). **c.** 1.5 x 10^9^ physical particles of LVTauS and LVTauAgg quantified by p24 ELISA were added to the original (TauRD-YT) and murinized (Tau-YTmCat1) reporter cells and induction of aggregation (FRET positive cells, maroon) and infection (mRuby positive cells, cyan) was quantified by flow cytometry. Mean ± SEM (n = 3 biological replicates). **d.** Optiprep gradient of sucrose cushion-pelleted LVTauAgg (detailed protocol in Supplementary Fig. 1c). Representative immunoblots of the Optiprep fractions are shown. Molecular weight (MW) standards are indicated. (n = 4 biological replicates). **e.** Induction of tau aggregation (seeding competence, FRET signal, cyan) and infection efficiency (mRuby signal, maroon) of the different Optiprep fractions from (d) quantified by flow cytometry using Tau-YTmCat1 cells. Mean ± SEM (n = 4 biological replicates).

To address the possibility that our findings may be limited to the isolated repeat domain of tau or the fluorescent tag YFP, we similarly produced LVs in HEK293T cells containing aggregated or soluble full-length tau (FLtau 0N2R, P301L/V331M) and tested seeding competence using the reporter cell line (Supplementary Fig. 1b). LVs produced in cells containing FLtau aggregates efficiently induced aggregation in the reporter cell line, while lentiviruses produced in cells containing soluble FLtau did not (Supplementary Fig. 1b), confirming that incorporation of tau aggregates into viral particles is not restricted to the repeat domain.

Extracellular vesicles are a byproduct of LV production. To discriminate between viruses and extracellular vesicles mediating aggregate seeding, we performed a previously established Optiprep (iodixanol) density gradient centrifugation to separate vesicles (light fractions) from viruses (heavy fractions)^26,32,33^ (Supplementary Fig. 1c). Interestingly, tau exclusively migrated in heavy fractions containing the viral capsid protein p24, the most abundant viral protein, indicating the presence of LV particles (Fig. 1d and Supplementary Fig. 1d). Next, we measured the infectivity and seeding competence of each Optiprep fraction. As expected based on p24 immunoblotting, the mature infectious viruses were in fractions 8 to 10, corresponding to the mRuby positive cell signal (Fig. 1d, e). Consistent with the presence of tau in the viral fractions, fractions 8 and 9, containing the mature infectious viral particles, showed the highest seeding efficiency, (Fig 1d, e). Of note, Optiprep did not enhance the seeding capacity of the particles, but rather at higher concentrations slightly decreased seeding with LVTauAgg (Supplementary Fig. 1e). We further purified the LV particles by pooling the fractions containing p24 and tau (F8-11) and pelleting them through a sucrose cushion (Supplementary Fig 1c). Purified LVTauAgg had properties consistent with an infectious virus, as it contained detectable levels of the viral protein p24 and was capable of inducing expression of the mRuby payload in infected cells (Supplementary Fig. 1f, g). Furthermore, purified LVTauAgg appeared to contain seeding-competent tau material, as it produced tau FRET signal in infected cells and contained tau protein, including the pathological marker pTau(Ser 356) (Supplementary Fig. 1f, g). The seeding competence of the purified particles was lower compared to particles prior to gradient purification when normalized to physical particle numbers (Fig. 1b and Supplementary Fig. 1g). This reduction is likely due to partial disruption of the viral envelope protein caused by mechanical forces during ultracentrifugation.

Taken together, these data indicate that tau aggregates can be incorporated into LV particles during production and that their presence seems to be compatible with the formation of mature, infectious virions. Moreover, viral particles loaded with aggregated tau induce tau aggregation in recipient cells in a virus receptor-dependent manner.

### Quantification and visualization of tau aggregates inside lentiviral particles

During viral production, tau aggregates could potentially associate with the outside of the viral particles (Fig. 1a and 2a). To determine whether tau was incorporated within the viral particles, we performed limited proteolysis using proteinase K (PK), a non-specific protease, in the presence or absence of detergent (Triton X-100). Quantification by immunoblotting revealed that ∼ 60% of tauRD-YFP was protected from proteolysis, indicating its location inside LV particles (Fig. 2a, b). LVs produced in cells containing soluble tau (LVTauS) also contained detectable amounts of tau, however these levels were markedly lower than those observed in LVTauAgg particles (Fig. 2a, c and Supplementary Fig. 2a). Notably, LV particles produced in naïve HEK293T cells did not become seeding-competent when incubated with lysates from cells containing aggregated tauRD-YFP (Supplementary Fig. 2b). This result suggests that, although tau aggregates may associate with the exterior of viral particles, such tau species are insufficient to induce aggregation in infected cells

**Figure 2.**
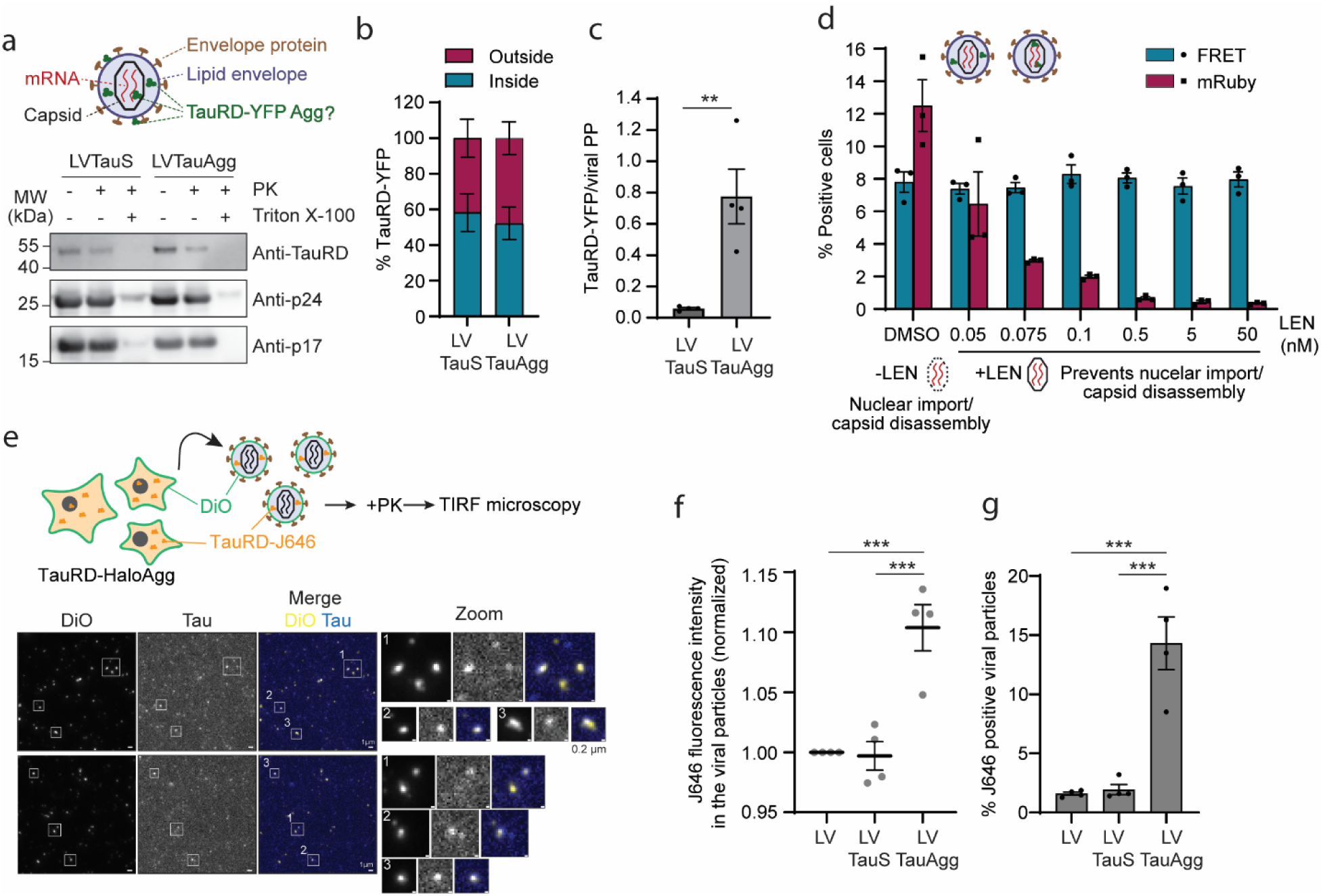
Quantification and visualization of tau aggregates inside viral particles. **a.** Simplified scheme of the structure of the LV indicating the possible locations of TauRD-YFP aggregates inside or outside the LV. LVTauS and LVTauAgg purified by Optiprep gradient (Supplementary Fig. 1c) were digested with proteinase K (PK) in the absence or presence of detergent (Triton X-100). Representative immunoblots are shown. Molecular weight (MW) standards are indicated. (n = 4 biological replicates). **b.** Quantification of TauRD-YFP normalized by p24 from immunoblots in (a). Inside: +PK bands; Outside: (Total, -PK band) - (Inside, +PK band). Mean ± SEM. **c.** Quantification of TauRD-YFP inside LVTauS and LVTauAgg particles. The number of TauRD-YFP molecules was calculated using the immunoblots shown in (a) and using purified TauRD-YFP as standard. The number of viral physical particles (PP) in the viral samples was quantified by ELISA of the p24 viral protein. Fold change of TauRD-YFP within the LVTauAgg particles compared to LVTauS is provided in Supplementary Fig. 2a. Mean ± SEM. **P < 0.01 by two-tailed Student’s t-test. **d.** Effect of the capsid inhibitor Lenacapavir (LEN) on LVTauAgg seeding and infection. LVTauAgg particles in the presence of increasing amounts of LEN were added to the murinized reporter cell line (Tau-YTmCat1) and the % fractions of FRET (induction of tau aggregation, cyan) and mRuby (infection, maroon) positive cells were quantified by flow cytometry. Mean ± SEM (n = 3 biological replicates). **e.** HEK293T cells containing soluble or aggregated TauRD-Halo labelled with Janelia Fluor 646 and lipid green dye DiO intercalated in the plasma membrane were used to produce LVs. These LV particles were then digested with proteinase K (PK) and visualized by TIRF microscopy. Naïve cells not containing TauRD-Halo but labelled with Janelia Fluor 646 and with lipid green DiO were used as control (LV). Representative microscopy pictures of LVTauAgg are shown. Scale bars, whole field images, 1 µm; Scale bars, zoom images, 0.2 µm. (n = 4 biological replicates). **f.** Quantification of tau intensity (J646 fluorescence intensity) in the LVs normalized by LV produced in naïve cells (representative microscopy images are shown in (e) and Supplementary Fig. 2c). Mean ± SEM. ***P < 0.001 according to one-way ANOVA with Tukey’s post hoc test. (n = 4 biological replicates). **g.** Percentage of viral particles containing tau (J646 fluorescence) in the LVs (representative microscopy pictures are shown in (e) and Supplementary Fig. 2c). Mean ± SEM. ***P < 0.001 according to one-way ANOVA with Tukey’s post hoc test. (n = 4 biological replicates).

We estimated the amount of tau incorporated into LVTauAgg particles by immunoblotting using purified recombinant tauRD-YFP protein as standard. In parallel, the number of viral particles in each preparation was quantified by ELISA detection of the p24 capsid protein. Based on these measurements, we estimated that around 8% of viral particles contain tau, assuming incorporation of small oligomeric species corresponding to around 10 mer. Given the limited internal volume of the viral particles (around 100-150 nm in diameter), we hypothesized that only small tau oligomers can be accommodated within the particles.

To investigate whether seeding competent tau species are located inside the viral capsid or between the capsid and the lipid envelope (Scheme in Fig. 2a), we used the capsid inhibitor Lenacapavir (LEN). After infection, the intact capsid is imported to the nucleus where it disassembles to allow integration of the viral genetic material. LEN stabilizes the capsid lattice and inhibits its nuclear import and disassembly^34,35^. As expected, cotreatment of the reporter cells with LVTauAgg and LEN resulted in a concentration dependent reduction of mRuby positive cells, confirming effective inhibition of viral capsid uncoating (Fig. 2d). In contrast, tau seeding competence remained unchanged, suggesting that seeding competent tau species are not located inside the viral capsid (Fig. 2d).

To visualize individual tau-containing viral particles, we used total internal reflection fluorescence (TIRF) microscopy, which provides high contrast with low background fluorescence. As in previous experiments, we used HEK293T cells containing either soluble or aggregated tau, however, in this case, tauRD was fused to Halo tag to enable labelling with the bright and stable fluorescent dye Janelia Fluor 646. To detect the viral particles, we used the lipid dye DiO, which intercalates in the cellular lipid bilayer. Therefore, tau-containing viral particles were labelled with DiO and Janelia Fluor 646 (Fig. 2e and Supplementary Fig. 2c). Prior to imaging, particles were digested with proteinase K to remove any potential tau bound to the outside of the particles. The tau signal was significantly higher in the LVTauAgg particles compared to LVTauS or LVs made in naïve HEK293T cells (Fig. 2e, f and Supplementary Fig. 2c). Around 14% of LVTauAgg particles contained tau (Fig. 2e, g and Supplementary Fig. 2c), consistent with the quantification upon PK digestion considering small tau oligomeric species of 10 mer (around 8%) (Fig. 2a).

In summary, tau aggregates reside somewhere outside the capsid in LV particles, with approximately 10% of particles containing tau oligomeric species.

### Gag polypeptide may be responsible for targeting tau aggregates into viral particles

To investigate how tau aggregates are incorporated into the LV particles, we first tested the interaction between tau and the viral polypeptide Gag by immunoprecipitation of tau from lysates of cells producing LVs containing soluble or aggregated tau, followed by mass spectrometry or immunoblotting (Fig. 3a, b). The Gag polypeptide was found to interact with both soluble and aggregated tau, exhibiting around ∼ 5-fold higher affinity for the aggregated species (Fig. 3a, b). Thus, incorporation of tau aggregates is likely driven by affinity for the viral Gag polypeptide.

**Figure 3.**
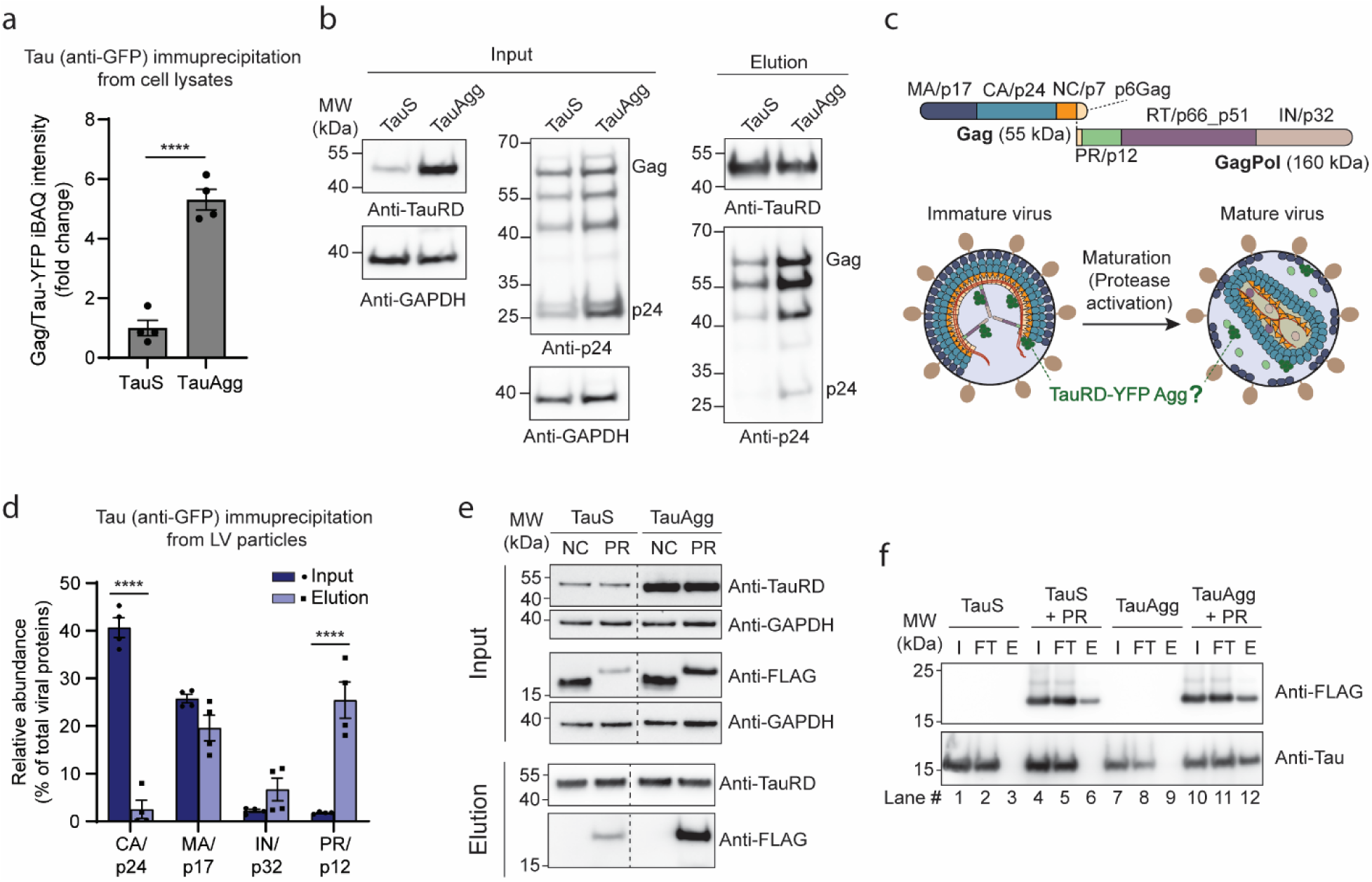
Tau aggregates interact with the viral Gag polypeptide and the viral protease. **a, b.** Interaction of Gag polypeptide with tau aggregates determined by tau (anti-GFP) immunoprecipitation of TauRD-YFPS (TauS) or TauRD-YFPAgg (TauAgg) cells producing LV particles, followed by mass spectrometry (a) or immunoblotting (b). Molecular weight (MW) standards are indicated. Mean ± SEM. ****P < 0.0001 by two-tailed Student’s t-test. (a, n = 4 biological replicates; b, n = 3 biological replicates). **c.** Simplified scheme of Gag and GagPol viral polypeptides (top) indicating the individual viral proteins and maturation of LV containing TauRD-YFP aggregates (bottom). **d.** Anti-GFP immunoprecipitation of tau aggregates from virus preparations followed by in-gel mass spectrometry. Tau aggregates inside the LVTauAgg particles were immunoprecipitated using anti-GFP beads. Input and elution fractions were separated on a SDS PAGE gel followed by in-gel mass spectrometry to differentiate the polypeptides from the individual viral proteins ((c) and Supplementary Fig. 3a). Relative abundance of each viral protein expressed as percentage of total viral proteins in input and elution fractions of the tau (anti-GFP) immunoprecipitation by mass spectrometry (calculated from iBAQ intensities). The highly abundant capsid (CA/p24) and matrix (MA/p17) viral proteins together with the viral integrase (IN/p32) and protease (PR/p12) are represented in the bar graph. Mean ± SEM. ****P < 0.0001 according to one-way ANOVA with Tukey’s post hoc test. (n = 4 biological replicates). **e.** Tau aggregates (anti-GFP) immunoprecipitation from cells expressing the nucleocapsid (NC) and PR viral proteins fused to 2xStrepTagII-3xFLAG tag. Representative immunoblots of input and elution fractions are shown. Molecular weight (MW) standards are indicated. (n = 3 biological replicates). **f.** Purified recombinant PR-FLAG was incubated with purified monomeric or aggregated tau *in vitro* for 30 min at 25 °C followed by anti-FLAG immunoprecipitation. Representative immunoblots are shown. Molecular weight (MW) standards are indicated. (n = 3 biological replicates).

LV particles are secreted as immature, non-infectious particles containing the Gag/GagPol polypeptides (Fig. 3c). Activation of the viral protease (PR) triggers cleavage of the Gag/GagPol polypeptide into individual viral proteins, facilitating the transition from immature to mature infectious viruses (Fig. 3c). To obtain further mechanistic insight into how tau aggregates are incorporated into the LV particles and their location therein, we immunoprecipitated tau aggregates from virus preparations, followed by in-gel mass spectrometry. This approach allowed us to distinguish individual viral proteins from the Gag/GagPol polypeptide (Fig. 3c and Supplementary Fig. 3a). The relative abundance of the viral protease (PR/p12) was highly increased in the aggregate eluate compared with the input (from ∼ 2% to 25%), suggesting an association with tau aggregates inside the viral particles (Fig. 3d). This interaction was confirmed by expression of the individual viral PR protein with a C-terminal FLAG tag in cells containing tau aggregates, followed by tau immunoprecipitation (Fig. 3e). To test the direct binding of PR to tau aggregates, we purified the viral PR with a C-terminal FLAG tag as well as recombinant tauRD (tau residues 244-371, C291A/P301L/C322A/V337M)^31^. PR-FLAG was incubated with monomeric or aggregated tau, followed by FLAG immunoprecipitation. PR specifically bound tau aggregates (lane 12) but not monomeric soluble tau (lane 6) (Fig 3f).

The viral nucleocapsid protein (NC/p7) that coats and condenses the genomic RNA inside the viral capsid was the only viral protein that was not detected in the immunoprecipitated by in-gel mass spectrometry (Fig. 3d and Supplementary Fig. 3b). Therefore, to probe for its binding to tau aggregates, we also expressed the viral NC fused to a FLAG tag in cells containing tau aggregates. Immunoprecipitation failed to show NC binding to tau aggregates (Fig. 3e), consistent with seeding remaining unaffected in the presence of the capsid stabilizer LEN (Fig. 2d). In agreement with these findings, LVTauAgg particles lacking mRNA payload, produced by expressing just the structural and envelop viral proteins, also incorporated tau aggregates. The resulting viral particles were seeding competent similar to those containing mRuby-tagged virus mRNA (Fig. 1d, e, Supplementary Fig. 1d, f and Supplementary Fig. 3c-f). These results indicate that tau aggregate incorporation and intraparticle localization are independent of elements residing within the viral capsid.

The Pol sequence within the GagPol viral polyprotein is not required for the assembly and release of viral particles. Therefore, to investigate the role of PR in viral incorporation of tau aggregates, we produced LVs in cells containing tau aggregates using plasmids in which either PR (delPR) or Pol (delPol) was deleted (Fig. 3c). PR and Pol proteins were not detected by mass spectrometry confirming the constructs (Supplementary Fig. 4a). When added to the reporter cells, both delPR and delPol LV particles were unable to induce aggregation (Fig. 4a). However, tau incorporation into the viruses was confirmed by both immunoblotting and mass spectrometry (Fig. 4b and Supplementary Fig. 4b), suggesting that PR and the whole Pol sequence are dispensable for tau packaging. To assess if tau contained within delPR and delPol LV particles was seeding competent, we first digested the particles with thermolabile proteinase K, followed by heat inactivation of the protease and lysis by sonication. After release of the aggregates from the particles, the lysates were added to the Tau-YTmCat1 reporter cells in the presence of the transfection reagent Lipofectamine (LPF), which is used to deliver protein aggregates into the cytoplasm of reporter cells^30,36–39^. As expected, tau species derived from delPR and delPol particles were seeding competent to a similar extent as those from WT particles (Fig. 4c). Notably, lysates from LVTauAgg particles exhibited higher seeding activity than intact LVs, likely indicating that during infection only a subset of LV particles successfully fuse with the plasma membrane and deliver tau species to the cytoplasm. These data indicate that, although tau aggregates can associate with viral PR, PR is not necessary for their incorporation into virions.

**Figure 4.**
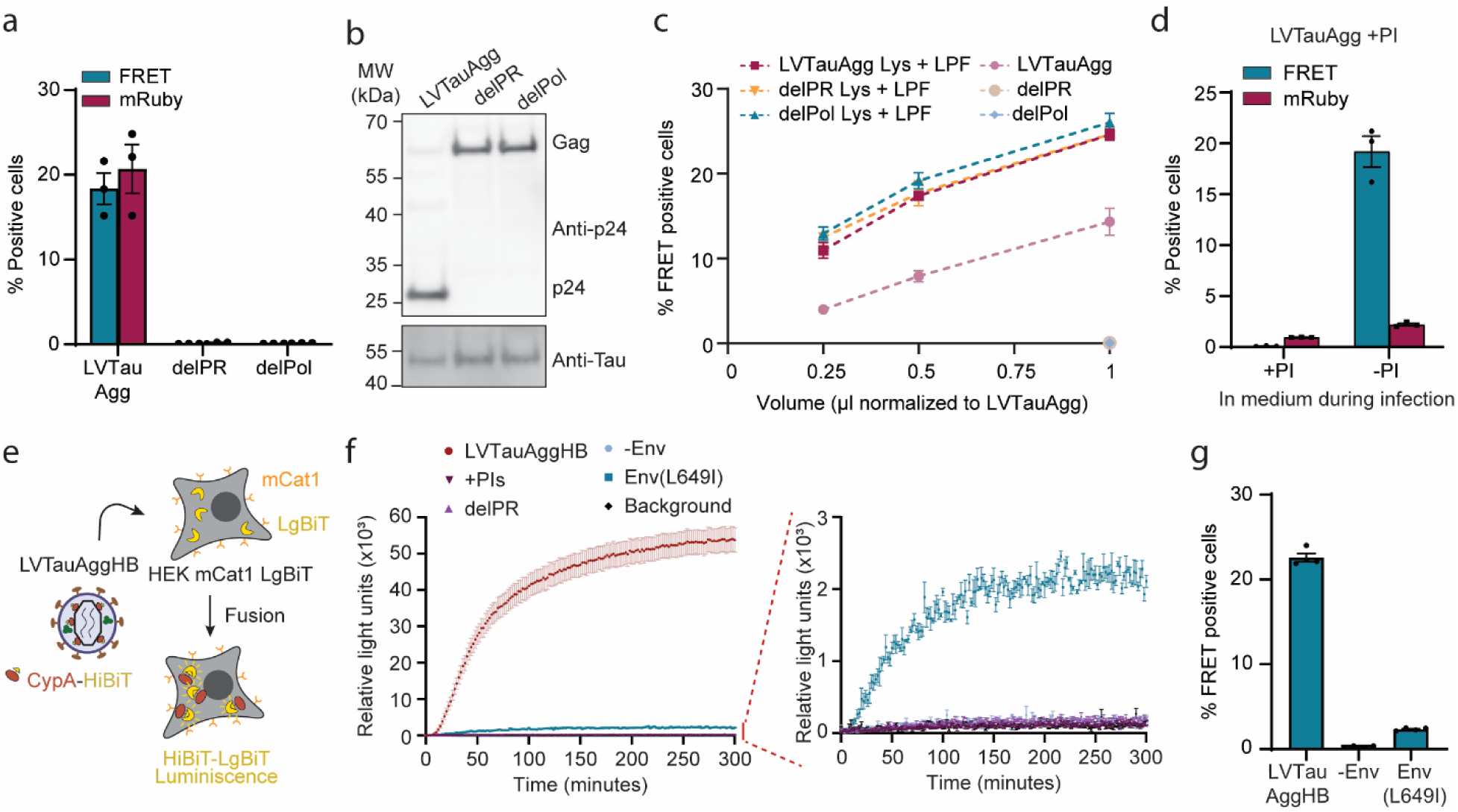
LV fusion with recipient cells and aggregate seeding is dependent on the viral protease and cleavage of the envelope protein. **a.** Seeding competence of LVTauAgg, delPR (LVTauAgg without viral PR) and delPol (LVTauAgg without the entire Pol sequence). Equivalent amounts of the LVTauAgg, delPR or delPol quantified by anti-p24 immunoblotting were added to the murinized tau seeding reporter cell line (Tau-YTmCat1). Induction of tau aggregation (FRET positive cells, cyan) and infectivity (mRuby positive cells, maroon) was quantified by flow cytometry. Mean ± SEM (n = 3 biological replicates). **b.** Representative immunoblots of LVTauAgg, delPR and delPol. Molecular weight (MW) standards are indicated. (n = 4 biological replicates). **c.** Seeding competence of LVTauAgg, delPR and delPol and their corresponding lysates (Lys). Equivalent amounts of the LV particles were digested with thermolabile proteinase K, followed by heat inactivation and addition of lipofectamine (LPF). The mixtures were then added to the murinized seeding reporter cell line (Tau-YTmCat1) and induction of tau aggregation (FRET positive cells) was quantified by flow cytometry. Mean ± SEM (n = 3 biological replicates, except for delPR Lys + LPF and delPR n = 4 biological replicates). **d.** LVTauAgg produced in the presence of protease inhibitors (PI) were added to the murinized tau seeding reporter cell line (Tau-YTmCat1) with PI present or absent in the culture medium during infection. Induction of tau aggregation (FRET positive cells, cyan) and infectivity (mRuby positive cells, maroon) was quantified by flow cytometry. Mean ± SEM (n = 3 biological replicates). **e.** Scheme of fusion assay. LVTauAggHB particles including the HiBiT subunit of the split nanoluciferase fused to cyclophilin A(CypA), a cellular host protein known to be incorporated into the viral particles, were added to the HEK293T reporter cell line expressing the viral receptor mCat1 together with the other half of the split nanoluciferase, the LgBiT subunit (HEK mCat1 LgBiT cells). After fusion, the two nanoluciferase subunits will interact and emit luminescence in the presence of substrate. **f.** LVTauAggHB, delPR (LVTauAggHB without viral PR), +PI (LVTauAggHB produced in the presence of PI), -Env (LVTauAggHB without envelope protein) and Env(L649I) (LVTauAggHB containing the uncleavable E-MLV env mutant, L649I) were added to the reporter cell line (HEK mCat1 LgBiT) and luminescence was monitored over time. The low luminescence signal is magnified in the graph on the right. Mean ± SEM (n=3 biological replicates, except for LVTauAggHB n = 11; delPR and +PI n= 4). **g.** Seeding competence of LVTauAggHB, -Env and Env(L649I). Equivalent amounts of the LVs normalized based on HiBiT incorporation were added to the murinized tau seeding reporter cell line (Tau-YTmCat1). Induction of tau aggregation (FRET positive cells) was quantified by flow cytometry. Mean ± SEM (n = 4 biological replicates).

To further explore the role of the viral PR protease activity in seeding, we produced LVs in cells containing tau aggregates in the presence of the PR protease inhibitors (PI), Indinavir and Ritonavir. These LV particles also incorporated tau (Supplementary Fig. 4b, c). When the PI were added to the medium during infection, the resultant LVs were unable to induce aggregation. However, when the PI were washed out by omitting them from the cell medium during infection, aggregate seeding competence was restored (Fig. 4d). Notably, the presence of inhibitors only during infection, and not during production of LVTauAgg particles, had no effect on infectivity or induction of aggregation (Supplementary Fig. 4d). Based on these data, tau seeding is specifically driven by particles containing viral proteins. Thus, the lack of seeding activity observed for LVTauAgg delPR, delPol and +PI particles is not due to impaired incorporation of tau aggregates into the viral particles, but more likely results from defective fusion of the viral particles with the plasma membrane.

### Fusion with recipient cells and aggregate seeding is dependent on viral protease

The E-MLV Env protein used to pseudotype the LVs is synthesized as a precursor protein that is cleaved in the secretory pathway into the surface subunit (SU) and the transmembrane subunit (TM). These two subunits remain covalently linked by a disulfide bond. SU-TM heterodimers assemble into trimers of dimers which are incorporated into the membrane of the budding virions. The 16 residue-long “R-peptide” located at the C-terminus of the TM domain, must be cleaved within the virions by PR to promote full maturation and acquisition of fusion capability (Supplementary Fig. 4e)^40^. Interestingly, mass spectrometry revealed a peptide containing the R-peptide cleavage site that was enriched in the seeding-inactive LVTauAgg delPR, delPol and +PI particles compared to the seeding-competent WT LVs (Supplementary Fig. 4e, f). We therefore hypothesized that viruses lacking active PR are unable to cleave the R-peptide and consequently fail to reach full fusion competence, rendering them incapable of inducing tau aggregation.

To assess this hypothesis, we used a split nanoluciferase system to detect viral fusion with the cell plasma membrane, in which half of the enzyme is incorporated in the viral particles (HiBiT subunit) and the complementary half is expressed in the reporter cell line (LgBiT subunit)^41,42^. Cells containing tau aggregates were transfected with the viral plasmids together with an additional plasmid encoding HiBiT-tagged cyclophilin A(CypA), a cellular protein known to be incorporated into the viral particles^41^, used here as a carrier to deliver HiBiT into the LVs. This resulted in the production of HiBiT-containing viral particles (LVTauAggHB). As a reporter cell line, we used HEK293T cells expressing the murine receptor mCat1 along with the LgBiT subunit (Fig. 4e and Supplementary Fig. 4g, h). We measured nanoluciferase luminescence over time, calculating the time required to reach half-maximum fusion (half-time) of viruses with the plasma membrane. Addition of WT particles to control HEK293T cells expressing only mCat1 or only LgBiT did not result in detectable luminescence (Supplementary Fig. 4g-i). Contrastingly, WT particles added to HEK mCat1 LgBiT cells rapidly fused to the plasma membrane with a half time of ∼60 min, as assessed by luminescence. As hypothesized, LV particles produced in the presence of the PI or lacking PR (delPR) were fusion incompetent (Fig. 4f and Supplementary Fig. 4j) and, similarly to previous results (Fig. 4a and d), failed to induce tau aggregation in the tau reporter cell line (Supplementary Fig. 4k).

To directly assess the contribution of the E-MLV Env R-peptide cleavage on fusion efficiency and seeding competence of the LV particles, we tested the fusion capacity of LVs produced without the E-MLV Env protein (-Env) or with the uncleavable L649I mutant^43^. As expected, LVs lacking the E-MLV Env protein did not fuse with the reporter cells and failed to induce aggregation in the Tau-YTmCat1 reporter cell line (Fig. 4f, g), confirming that the seeding capacity of LVTauAgg is receptor-dependent (Fig. 1c). The presence of tau inside LVTauAggHB -Env and Env(L649I) was confirmed by immunoblotting (Supplementary Fig. 4l). The Env(L649I) mutant showed a ∼95 % reduced fusion capacity compared to WT LV particles (LVTauAggHB), in line with a 90% reduction in seeding capacity (Fig. 4f, g and Supplementary fig. 4j). Therefore, a major loss of seeding competence in LVTauAgg produced with inactive PR is attributable to impaired fusion resulted from uncleaved E-MLV Env protein, rather than to a global maturation defect. Accordingly, the vast majority of LVTauAgg seeding competence derives from particles containing active PR.

Thus, while tau aggregates may bind the viral PR within the viral particles, neither PR nor the full Pol sequence is required for the incorporation of aggregates into virions. Instead, virus PR activity is essential for acquiring fusion efficiency through E-MLV Env R-peptide cleavage and subsequent delivery of tau seeds into the cytoplasm of the infected cells.

### Virus-mediated induction of tau aggregation as a tool in models of neurodegeneration

The development of appropriate human cellular models to study protein aggregation and spreading in neurodegeneration is urgently needed to understand the molecular mechanisms underlying this process and identify novel therapeutic targets. Despite their amyloidogenic nature, proteins like tau are remarkably resistant to aggregation in cellular models. As mentioned above, a widely used technique to induce aggregation in cells is to produce aggregates *in vitro* (preformed fibrils, PFFs) from recombinant purified protein and transfect those aggregates into cells using the classical lipid-based DNA transfection reagent LPF^30,36–39^. However, transfection reagents like LPF perturb the endolysosomal system and are toxic for some specialized postmitotic cells such as neurons^44–46^. Thus, the virus-mediated tau aggregate induction methodology shown here may be an efficient and more versatile (receptor specific) alternative to induce aggregation in different cellular models.

We assessed endolysosomal integrity upon aggregate delivery by LV particles or LPF mediated transfection (using PFFs or lysate from aggregate-containing cells) using a HEK293T Galectin 3 (Gal3) FRET (mRuby-Gal3 and mClover-Gal3, RC-Gal3) reporter cell line. Upon rupture of endolysosomal membranes, inner glycans become exposed and recruit clusters of Gal3 molecules, resulting in FRET signal (Fig. 5a). For direct comparison of tau aggregate seeding efficiency, we additionally performed the same experiment with Tau-YTmCat1 cells reporting aggregate formation (Fig. 1a). As anticipated, the use of LPF alone or combined with tau PFFs (or aggregate-containing cell lysate), generated Gal3-FRET positive cells, indicating impairment of endolysosomal integrity (Fig. 5b). To test the effect of the LV particles, we first incorporated the viral receptor mCat1 into the reporter cell membrane using exosome-like particles coated with the receptor allowing for efficient viral infection. After 2 hours, the LVTauAgg or LVTauS were added and FRET reporter signal recorded after a further 24 hours (Fig. 5a, b). Notably, the addition of LV particles did not damage the endolysosomal pathway (Fig. 5b), despite inducing tau aggregation (in the case of LVTauAgg) with similar efficiency as samples transfected with LPF and PFFs or aggregate-containing cell lysate (Fig. 5c).

**Figure 5.**
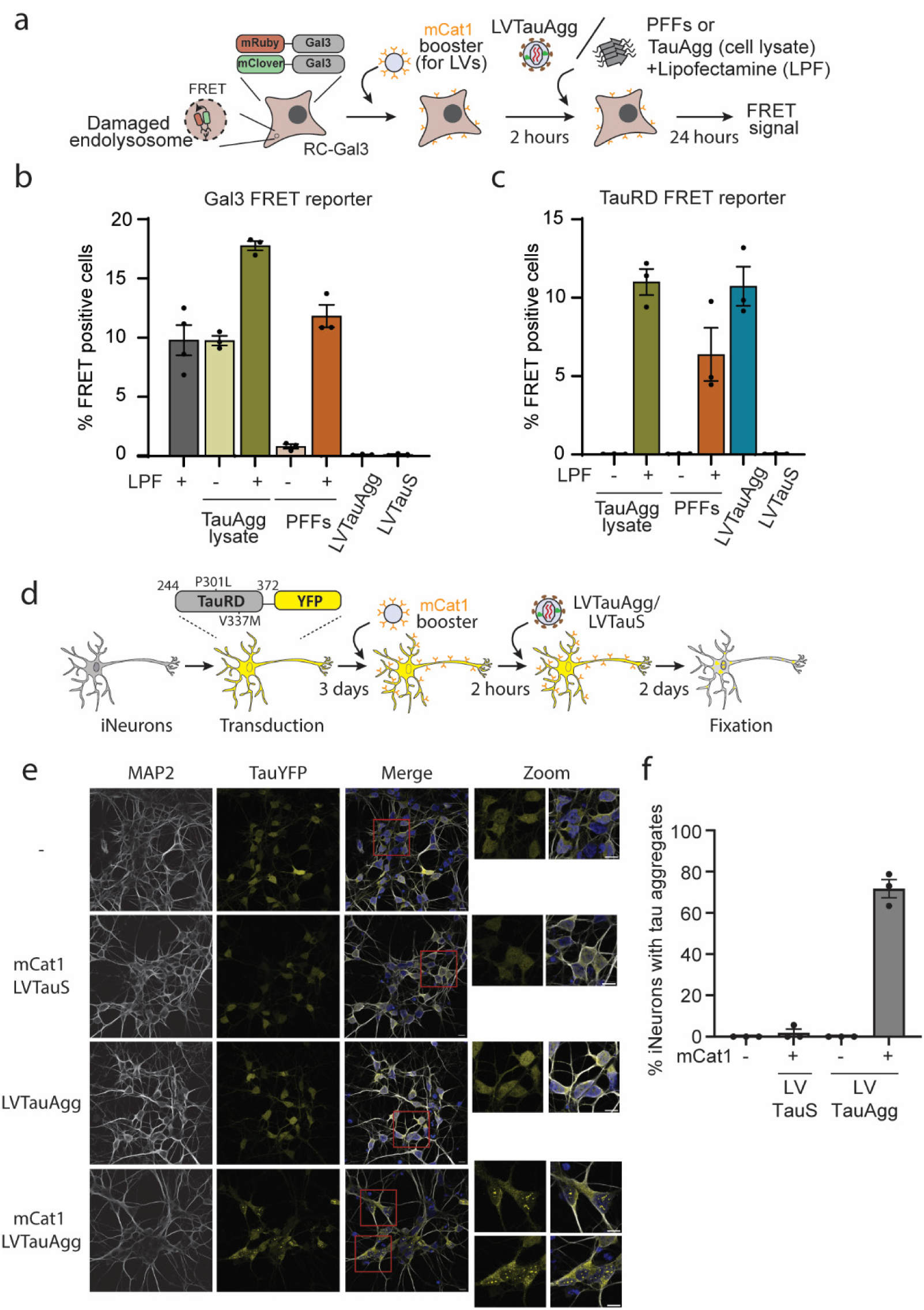
Virus-mediated induction of tau aggregation in human iNeurons. **a.** Schematic depicting the experimental strategy to assess effects of lipofectamine (LPF) and LV particles on endolysosomal integrity. Exosome-like particles coated with the viral receptor mCat1 (mCat1 booster, Takara) were added to the cells to allow subsequent LV infection. After 2 hours, LVTauAgg or LVTauS particles, LPF alone or mixed with tau preformed fibrils (PFFs, aggregates produced *in vitro* with purified recombinant TauRD) or aggregate-containing cell lysate, were added to the reporter cell line HEK293T expressing Galectin 3 (Gal3) FRET constructs (mRuby-Gal3 and mClover-Gal3, RC-Gal3). **b.** LPF alone or mixed with aggregate-containing cell lysate or tau PFFs, as well as LVTauAgg and LVTauS particles were added to the RC-Gal3 reporter cell line. For addition of LV particles, cells were pretreated with mCat1 booster (a). Induction of endolysosomal damage (FRET positive cells) was quantified by flow cytometry. Mean ± SEM (n = 3 biological replicates, except for LPF+ n= 4 biological replicates). **c.** LPF mixed with aggregate-containing cell lysate or tau PFFs, as well as LVTauAgg and LVTauS particles were added to the TauRD-YTmCat1 reporter cells (Fig. 1a). Induction of tau aggregation (FRET positive cells) was quantified by flow cytometry. Mean ± SEM (n = 3 biological replicates). **d.** Workflow for LVTauAgg seeding experiments in human induced pluripotent stem cells (iPSCs)-derived cortical neurons (iNeurons). iNeurons were transduced to express TauRD(P301L/V337M)-YFP and mCat1 was transiently induced to allow infection as in (a) using exosome-like particles coated with the viral receptor mCat1. Two hours later, LVTauAgg or LVTauS were added to the cultures and tau-YFP aggregation was assessed by fluorescence microscopy after further 48 hours. **e.** Representative fluorescence microscopy images of iNeurons expressing TauRD-YFP (yellow) incubated with LVTauAgg or LVTauS in the absence or presence of the viral receptor mCat1. iNeurons were stained with an antibody against the neuronal marker MAP2 (white). DAPI nuclear staining signal is additionally shown in blue in the merge. Enlarged sections are highlighted with red squares. Scale bars, 10µm. (n = 3 biological replicates). **f.** Quantification of iNeurons containing TauRD-YFP aggregates from fluorescence microscopy images from (e). Mean ± SEM (n = 3 biological replicates).

To directly determine the seeding efficiency relative to the amount of delivered tau aggregate (specific seeding efficiency), we estimated the amount of tau in the lysate of cells with tau aggregates by quantitative immunoblotting (Supplementary Fig. 4m) and compared it to the amount of tau within virus particles (Fig. 2a, b) and tau PFFs. When normalized to the same amount of tau, LVTauAgg were approximately 6,000-fold more efficient at inducing tau aggregation than *in vitro* generated tau PFFs delivered with LPF and ∼ 23-fold more efficient than tau aggregates derived from cell lysates delivered with LPF. Thus, delivery of tau aggregates within LV particles is markedly more efficient and less disruptive than LPF-mediated transfection.

Next, we evaluated virus-mediated tau aggregation in human induced pluripotent stem cells (iPSCs)-derived cortical neurons (iNeurons), a cell type for which LPF is inefficient and toxic^44–46^. We transduced iNeurons to express tauRD-YFP and introduced the viral receptor mCat1 using coated exosome-like particles, as for the RC-Gal3 reporter cell line (Fig. 5a, d). Two hours later, pseudotyped LV particles produced in cells containing either soluble or aggregated tau were added to the cultures. After 48 hours, the formation of tauRD-YFP aggregates was assessed by fluorescence microscopy (Fig. 5e, f). LVTauAgg rapidly and efficiently induced tau aggregation in up to 70% of iNeurons, but only when the receptor was present (Fig. 5e, f). In contrast, LVTauS failed to induce aggregation in iNeurons, even in presence of the receptor (Fig. 5e, f).

In summary, pseudotyped LVs are a highly efficient tool for inducing aggregation in LPF sensitive cells, such iNeurons. Moreover, owing to their selectivity for the mCat1 receptor, LVs can be used to target specific subpopulations of cells.

## Discussion

Transcellular propagation of protein aggregates, a likely key factor in the progression of major NDs^1,3,47^, can occur via different mechanisms, including transfer via extracellular vesicles^5,6,22^. Interestingly, extracellular vesicles closely resemble enveloped viruses in both structure and function, as both are surrounded by a lipid bilayer and serve as carriers of nucleic acids and proteins that can be transferred to recipient cells following membrane fusion^26,27^. A key distinction, however, lies in membrane fusion efficiency. Viruses have evolved highly optimized, rapid, and tightly regulated entry mechanisms^48^, whereas extracellular vesicles exhibit substantially lower fusion efficiency due to the absence of specialized fusion machinery. Moreover, escape of extracellular vesicle cargo from endosomal compartments is inefficient, resulting in degradation rather than delivery to the cytosol^49^.

Here we show that LV production in cells containing tau aggregates results in the incorporation of seeding competent tau species into viral particles that can template tau aggregation in infected cells in a viral receptor-dependent manner (Fig. 1a, c and Fig. 4g). Our biochemical characterization revealed efficient incorporation of tau aggregates into LV particles, with ∼10-15 % of particles containing aggregate material (Fig. 2a, g). The conformation of these tau seeds remains unknown, but given the restricted space available within the viral particles (around 100-150 nm diameter), incorporated tau species are likely small oligomers. This would be consistent with reports that very small tau oligomers and even monomeric tau in a specific conformation may be seeding-competent^38,50,51^.

Our findings are consistent with previous reports demonstrating efficient intercellular induction of tau aggregation in coculture systems in which donor cells co-express the viral envelope protein together with Gag/GagPol polypeptide^25^. However, in contrast to these experiments, where envelope protein expression alone appears sufficient to promote aggregation, in our system using isolated particles, viral protease (PR) activity is required to confer seeding competence to the LV particles. Approximately 90% of the dependence of LV-mediated tau seeding on PR activity can be attributed to the essential role of proteolytic E-MLV Env processing in establishing fusion competence^40^ (Fig. 4e-g and Supplementary Fig. 4j). This, together with the observation that tau aggregates were detected almost exclusively in density gradient fractions containing infectious viruses (Fig. 1d, e), supports the conclusion that tau aggregate seeds can be packaged inside mature viral particles.

The E-MLV Env protein, a member of the gamma-type envelope family, possesses a C-terminal R-peptide present in the immature envelope trimer that the viral protease must cleave to enable fusion competence^40^(Supplementary Fig. 4e). Accordingly, LV particles lacking protease or harbouring an inactive protease displayed a marked accumulation of uncleaved R-peptide (Supplementary Fig. 4e, f). These particles, as well as LVs containing the uncleavable E-MLV Env mutant L649I, were essentially deficient in membrane fusion and tau seeding (Fig. 4e-g and Supplementary Fig. 4j, k). Notably, washout of protease inhibitors rescues seeding competence (Fig. 4d), demonstrating that seeding activity arises almost exclusively from protease-active, fusion-competent viral particles. Consistent with this hypothesis, extracellular vesicles (EVs) decorated with certain non-gamma-type viral envelope proteins that do not require proteolytic processing to acquire fusion competence, such as the SARS-CoV-2 spike S and the vesicular stomatitis virus glycoprotein^40,41,52,53^, have been reported to deliver proteopathic seeds to the cytoplasm of recipient cells more efficiently than uncoated EVs^24^.

Viruses incorporate some host cellular proteins, either via a non-specific passive process or direct, affinity-driven interactions between viral and host proteins^28,29^. Although the viral PR specifically bind tau aggregates (Fig. 3d-f), neither its presence nor any other component of Pol is required for the incorporation of tau seeding-competent species into LV particles (Fig. 4b, c and Supplementary Fig. 4b). Moreover, none of the elements located within the viral capsid appear to facilitate incorporation (Fig. 2d, 3e and Supplementary Fig. 3c-f). In contrast, affinity of the viral structural polyprotein Gag for tau aggregates (Fig. 3a, b) may mediate the incorporation of tau seeds into LV particles. Further studies are required to elucidate the specific elements within the Gag polyprotein, or additional shared binding partners, responsible for the incorporation of tau aggregates into LV particles and to determine whether this phenomenon also occurs with other enveloped viruses.

Epidemiological and experimental evidence accumulated over the past three decades implicates viral infection in ND pathogenesis. Nevertheless, this area of research remains relatively underexplored, and the contribution of viruses to NDs remains controversial^12–21^. In addition to classical viral infections that can reach the brain and possibly persist during aging, such as HIV or HSV-1, both of which have been linked to neurodegeneration^16,54^, it has recently been proposed that endogenous viral elements may also contribute to disease pathogenesis. Human endogenous retroviruses (HERVs) and the neuronal activity-regulated protein Arc have emerged as potential contributors to disease progression by facilitating the transcellular spread of pathological protein aggregates^25,55,56^. Notably, Arc contains a retroviral Gag-like domain that enables the formation of extracellular vesicle-associated capsids and has been reported to interact with tau^56^. The human genome contains multiple HERV loci, many of which still retain gamma-type envelope open reading frames that require protease-mediated cleavage^40^, similar to the LV mechanism described here. Importantly, accumulating evidence indicates that HERVs reactivate during aging and in NDs^57^.

Our data support and expand the hypothesis that viruses and endogenous virus-like particles may incorporate tau aggregates and serve as highly efficient vehicles for cytoplasmic delivery in infected cells, possibly contributing to the transcellular propagation of proteopathic tau species. In this context, our results suggest that antiviral interventions directed to viral proteases or envelope proteins could potentially mitigate disease progression by restricting fusion-competent particle formation.

Understanding the molecular mechanisms that drive ND pathogenesis and progression is essential for the development of effective therapies. However, modelling human neurodegeneration has long been challenging, a difficulty reflected in the lack of cost-effective treatments and the high failure rate of clinical trials^58^. Recent advances in stem-cell-based *in vitro* systems, including organoids and other three-dimensional (3D) cellular models which recapitulate key disease hallmarks like amyloid beta aggregation and tau phosphorylation, have opened new opportunities^59–61^. Despite this progress, the spontaneous formation of tau aggregates, as observed in patient brains, or their timely induction through the addition of exogenous PFFs remains difficult in these models^60–63^. Consequently, studying the mechanisms underlying transcellular propagation of aggregates is still limited. LVs present a versatile tool enabling the rapid and efficient induction of tau aggregation. LVs containing tau aggregates are approximately 20-fold more efficient at inducing tau aggregation in recipient cells and less disruptive of the endolysosomal pathway than comparable amounts of tau aggregates derived from the same cell lysates delivered via LPF (Fig. 5a-c). Notably, aggregation is achieved in 70% of iNeurons just 48 hours after infection (Fig. 5d-f). In addition, the strict specificity of E-MLV Env pseudotyped viruses for murine cells (Fig. 1a, c) confers enhanced biosafety and practical advantages compared to human-tropic viral systems^64^. Importantly, this specificity of tau-containing pseudotyped viruses for the mCat1 receptor provides a unique opportunity to selectively induce tau aggregation in defined cell types or restricted subsets of cells. This can be achieved, for example, through cell-type specific promotors or transient receptor expression prior to the assembly of multicellular 3D models. Moreover, the requirement for active viral PR to acquire tau seeding-competence, together with the observation that some PI appear to be reversible (Fig. 3h), adds an additional layer of fine-tuned control to the system. Such an approach would enable precise spatial control of aggregation initiation and allows subsequent investigation of tau spreading and seeding mechanisms in neighbouring cells. This platform, combined with recent advances in generating disease-relevant tau fibrils in vitro^65,66^, has the potential to substantially accelerate our understanding of ND mechanisms and therefore the development of novel therapeutic strategies.

## Methods

### Plasmids

The psPAX2 packing plasmid used for LVs production, was a gift from Didier Trono (Addgene plasmid # 12260; http://n2t.net/addgene:12260; RRID:Addgene_12260). The deletions in the psPAX2 plasmid, psPAX2delPR and psPAX2delPol were synthesized (Twist Bioscience) and cloned in the previously digested psPAX2 plasmid with EcoRI (New England Biolabs) followed by assembly using the NEBuilder HiFi DNA assembly master mix (New England Biolabs). The pMD2.G-E-MLV Env plasmid used for LVs production was obtained by PCR amplification of the backbone from the pMD2.G plasmid, gift from Didier Trono (Addgene plasmid # 12259; http://n2t.net/addgene:12259; RRID:Addgene_12259) and the ecotropic Murine Leukemia Virus envelope protein from the pHCMV-EcoEnv plasmid, gift from Miguel Sena-Esteves (Addgene plasmid # 15802; http://n2t.net/addgene:15802; RRID:Addgene_15802)^67^ followed by assembly using the NEBuilder HiFi DNA assembly master mix (New England Biolabs). The pMD2.G-E-MLV Env(L649I) was obtained by PCR amplification followed by assembly using the NEBuilder HiFi DNA assembly master mix (New England Biolabs). The pLenti-mRuby3 plasmid and pLenti-mClover3 plasmids used for LVs production were obtained by PCR amplification of mRuby3 and mClover3 from the pKanCMV-mClover3-mRuby3 plasmid, gift from Michael Lin (Addgene plasmid # 74252; http://n2t.net/addgene:74252; RRID:Addgene_74252)^68^ and the lenti-EF1a-dCas9-VPR-Puro plasmid, gift from Kristen Brennand (Addgene plasmid # 99373; http://n2t.net/addgene:99373; RRID:Addgene_99373)^69^ followed by assembly using the NEBuilder HiFi DNA assembly master mix (New England Biolabs). The pLenti-mCat1 plasmid used for murinizing mammalian cells was obtained by mCat1 PCR amplification from the plasmid pp-mCAT1, gift from Vincent Lotteau & Philippe Mangeot (Addgene plasmid # 83948; http://n2t.net/addgene:83948; RRID:Addgene_83948)^70^ and lenti-EF1a-dCas9-VPR-Puro plasmid, followed by assembly using the NEBuilder HiFi DNA assembly master mix (New England Biolabs). CMV-LgBit plasmid was a gift from Nevin Lambert (Addgene plasmid # 234738; http://n2t.net/addgene:234738; RRID:Addgene_234738)^71^. The pN1-CypAHiBiT was obtained by cloning the CypAHiBit ordered as a gene fragment (Twist Bioscience) into the mTurquoise2-N1 backbone, gift from Michael Davidson & Dorus Gadella (Addgene plasmid # 54843; http://n2t.net/addgene:54843; RRID:Addgene_54843)^72^ digested with BamHI and NotI restriction enzymes, followed by assembly using the NEBuilder HiFi DNA assembly master mix (New England Biolabs). The TauRD(P301L/V337M)-Halotag sequence was cloned in the backbone of the pCW57.1-MAT2A plasmid which was a gift from David Sabatini (Addgene plasmid # 100521; http://n2t.net/addgene:100521; RRID:Addgene_100521)^73^ to generate the pCW-TetOffTauRD(P301L/V337M)-Halotag plasmid. Gal3-mRuby3-IRES-Gal3-mClover3 was cloned in the backbone of Plasmid EGFP-α-SynA53T (Addgene plasmid #40823, gift from David Rubinsztein)^74^. To generate the pLenti-PR-FLAG and pLenti-NC-FLAG plasmids, codon optimized (GenScript) versions of PR and NC proteins with C-terminal 2xStrepTagII-TEVS-3xFLAG for mammalian expression similar to previously described^75^ were synthesized (Twist Bioscience) and cloned in the lenti-EF1a-dCas9-VPR-Puro plasmid for mammalian expression and in a pHUE^76^ for *E. coli* expression and purification (pHUE-PR(D25N)-FLAG plasmid). The catalytic site of PR was inactivated by introducing the D25N mutation to avoid cellular toxicity.

PR(D25N) 2xStrepTagII-TEVS-3xFLAG amino acid sequence:

MPQITLWQRPLVTIKIGGQLKEALL**N**TGADDTVLEEMNLPGRWKPKMIGGIGGFIKV RQYDQILIEICGHKAIGTVLVGPTPVNIIGRNLLTQIGCTLNF<u>GAAAGWSHPQFEKGGG</u> <u>SGGGSGGGSWSHPQFEKGENLYFQGADYKDHDGDYKDHDIDYKDDDDK</u>

NC 2xStrepTagII-TEVS-3xFLAG amino acid sequence:

MQRGNFRNQRKIVKCFNCGKEGHTARNCRAPRKKGCWKCGKEGHQMKDCTERQANGAAAGWSHPQFEKGGGSGGGSGGGSWSHPQFEKGENLYFQGADYKDHDGDYKDHDIDYKDDDDK

### Cell lines

HEK293T cells expressing tauRD (P301L/V337M)-YFP in soluble (TauRD-YFPS cell line) or aggregated (TauRD-YFPAgg cell line) form, FLtau(P301L/V337M) in soluble (FLTauS cell line) or aggregated (FLTauAgg cell line) form, the HEK293T FRET reporter cell line expressing tauRD (P301L/V337M) N-terminally fused to either EYFP or mTurquoise2 (TauRD-YT cell line) were previously described^30,31,77^.

HEK293T cells for lentiviral packaging (Lenti-X 293T cell line, Takara) expressing tauRD (P301L/V337M) N-terminally fused to Halotag (TauRD-Halo cell line) and HEK293T TauRD-YT cells expressing mCat1(Tau-YTmCat1 cell line) were generated by transduction with the corresponding lentiviruses (see “Lentiviral production”) followed by antibiotic selection. Monoclonal cell lines were then selected by flow cytometry. To generate the TauRD-Halo tag cell line stably propagating tau aggregates (TauRD-HaloAgg), TauRD-Halo cells were transfected with 1 µg total protein from TauRD-YFPAgg cell lysate using lipofectamine 3000 (Thermo Fisher Scientific) similar to that describe below (see “Induction of tau aggregation in cells”). TauRD-YFPAgg cells were lysed in 1% Triton X-100/PBS (Thermo Fisher Scientific) supplemented with protease inhibitor cocktail (MERCK, EDTA-free) and benzonase (prepared in-house) and kept on ice for 20 min. Protein concentration was quantified using BCA Protein Assay (Thermo Fisher Scientific). A monoclonal cell line stably propagating aggregates was selected by addition of Janelia Fluor 549 Halo tag ligand (Promega), sorting in a 96 well plate and fluorescent microscopy screening.

HEK293T cells stably expressing PR(D25N), NC, LgBiT or mRuby3-Galectin 3 and mClover3-Galectin 3 (RC-Gal3 cell line) were generated by transfection of the pLenti-PR-FLAG, pLenti-NC-FLAG, CMV-LgBit or Gal3-mRuby3-IRES-Gal3-mClover3 plasmids respectively using lipofectamine 3000 (Thermo Fisher Scientific) following manufacturer’s instructions and subsequent antibiotic selection.

HEK293T cell lines were maintained at 37 °C and 5% CO_2_ in Dulbecco’s modified Eagle’s medium (DMEM High Glucose, L-glutamine, Phenol Red, Sodium pyruvate, Thermo Fischer Scientific) supplemented with 10% fetal bovine serum (Thermo Fisher Scientific), 100 U mL^−1^ penicillin (Thermo Fisher Scientific), 100 µI mL^−1^ streptomycin sulfate (Thermo Fisher Scientific). TauRD-YT and TauRD-YTmCat1 cell lines were maintained in the medium described above supplemented with 200 µg mL^−1^ G418 (Thermo Fisher Scientific).

iPSCs line HPSI0214i-kucg_2 (RRID:CVCL_AE60) was purchased from the UK Health Security Agency (77650065, supplied by HipSci). iPSCs were maintained at 37 °C and 5% CO2 in mTeSR or mTeSR plus medium (Stem Cell Technologies) on Geltrex-coated (Thermo Fisher Scientific) cell culture plates. Cells were split when confluent using ReLeSR (Stem Cell Technologies).

Neural progenitor cells (NPCs) and iPSCs-derived forebrain-type neurons were generated as described previously^78^ using the STEMdif SMADi neural induction kit (Stem Cell Technologies) following the monolayer protocol, the STEMdiff Forebrain Neuron Differentiation Kit (Stem Cell Technologies) and the STEMdiff Forebrain Neuron Maturation Kit (Stem Cell Technologies). A detailed differentiation protocol is available from protocols.io (https://doi.org/10.17504/protocols.io.eq2lyxz3wgx9/v1).

### Induction of tau aggregation in cells

Tau seeding in HEK293T reporter cells: 110,000 cells per well of the HEK293T reporter cell line Tau-YTmCat1 or Tau-YT were dispensed into a 12-well plate. 16-24 h later, LV particles were directly added to the cells. Comparable amounts of different LVs were added to the cells by normalization using HIV Type 1 p24 antigen ELISA 2.0 (ZeptoMetrix, Biozol), anti-p24 immunoblotting (when PR was deleted or inhibited because the Gag polypeptide was not detected by p24 ELISA) or HiBiT incorporation using the Nano-Glo HiBiT Lytic Detection System (Promega).

When quantifying the seeding capacity of tau aggregates inside the LV particles, similar amounts of LV particles normalized by p24 immunoblotting were digested with thermolabile proteinase K (NEB) for 30 min at 22 °C. Proteinase K was then inactivated by incubation for 10 min at 55 °C. The samples were sonicated using a Bioruptor sonication bath (Diagenode) (25 cycles of 5 s on – 5 s off) and different volumes were transfected using lipofectamine 3000 (Thermo Fisher Scientific) as described below.

When comparing seeding competence of tau aggregates delivered by LV particles or LPF mediated transfection, 2.8×10^8^ physical particles of LVTauS or LVTauAgg quantified by HIV Type 1 p24 antigen ELISA 2.0 (ZeptoMetrix, Biozol) were added to the wells. When seeding with tau PFFs or aggregate-containing cell lysate, tau aggregates were transfected with lipofectamine 3000 (Thermo Fisher Scientific). Cell pellets were lysed in 1% Triton X-100/PBS (Thermo Fisher Scientific) supplemented with protease inhibitor cocktail (MERCK, EDTA-free) and benzonase (prepared in-house) and kept on ice for 20 min. Protein concentration was quantified using BCA Protein Assay (Thermo Fisher Scientific). Specifically, 55 ng of tau PFFs (see “Protein expression, purification and aggregation”) or 0.2 µg total protein from tau aggregate-containing cell lysate were mixed with a mixture of 50 µL Opti-MEM (Thermo Fisher Scientific) and 1.6 µl lipofectamine 3000 reagent (Thermo Fisher Scientific) and incubated for 20 min at room temperature. The mixtures were then added to the cells.

When naïve LVs (produced in the lentiviral packaging Lenti-X 293T cell line, Takara) were incubated with aggregate-containing cell lysate, TauRD-YFPAgg cell pellets were resuspended in PBS supplemented with protease inhibitor cocktail (MERCK, EDTA-free) and benzonase (prepared in-house) and sonicated using a Bioruptor sonication bath (Diagenode) (30 cycles of 30 s on – 30 s off). Total protein was quantified using the Pierce Dilution-Free Rapid Gold BCA Protein Assay (Thermo Fisher Scientific). Naïve LVs were incubated with 1 µg total protein in a total volume of 20 µL for 2 hr at 37 °C and then added to the reporter cells. 1 µg total protein was directly added to the cells or incubated with 50 µL Opti-MEM (Thermo Fisher Scientific) and 1.6 µl lipofectamine 3000 reagent (Thermo Fisher Scientific) for 20 min at room temperature. The mixture was then added to the cells.

When adding Lenacapavir (MedChemExpress) the corresponding concentrations were directly added to the medium of the reporter cells.

16–24 h (when using LV particles without viral mRNA or aggregates transfected with lipofectamine) or 48 h (when using LV particles containing expressing RNA) after seeding, cells were collected using TrypL Express Enzyme (Thermo Fisher Scientific), fixed with 4% paraformaldehyde (PFA) in PBS for 10 min, washed with PBS and resuspended in PBS for analysis with an Attune NxT flow cytometer using the Attune Cytometric software Attune NxT Software v. 5.1.1 or v.6.2.3337.1 (Thermo Fisher Scientific). To measure the mTurquoise2 and FRET fluorescence signals, cells were excited with 405 nm laser light and fluorescence was determined using 440/50 and 530/30 filters, respectively. To measure the YFP fluorescence signal, cells were excited at 488 nm and emission was recorded using a 530/30 filter. Data processing was performed using FlowJo V10 and V10.10.0 software (FlowJo LLC). After gating single cells, an additional gate was introduced to exclude YFP-only cells that show a false-positive signal in the FRET channel due to excitation at 405 nm. The FRET positive gate was set by plotting the FRET fluorescence signal versus the mTurquoise2 fluorescence signal using as reference non-seeded cells. Detailed gating strategy in^31^.

Tau seeding in iNeurons: 100,000 iNeurons cultured in a well of a 24-well plate were transduced with tauRD (P301L/V337M)-YFP LVs (see “Lentivirus production”). After 72 hr, 2.5 µL of the ecotropic booster receptor (Takara) was added to the cells and the plate was centrifuged at 1,200 x g for 15 min at 18 °C and placed back in the incubator. After 2 hr, 0.7×10^9^ physical particles of LVTauS or LVTauAgg quantified by HIV Type 1 p24 antigen ELISA 2.0 (ZeptoMetrix, Biozol) was added to the wells and the plate was centrifuged at 1,200 x g for 15 min at 18 °C and placed back in the incubator. After 48 hr, the cells were washed with PBS and fixed with 4% PFA/PBS for 10 min, washed with PBS and permeabilized with 0.1% Triton-X100/PBS for 5 min. Blocking solution (8% BSA/PBS) was added for 1 hr. Coverslips were then transferred to a humid chamber and incubated overnight with anti-MAP2 antibody (PA1-10005, Thermo Fisher Scientific) diluted in 1% BSA/PBS. Cells were washed with PBS, and incubated with goat anti-chicken IgY (H+L), Alexa Fluor 647 (A-21449, Thermo Fisher Scientific, 1/500 dilution) diluted in 1% BSA/PBS for 1 hr, washed with PBS and stained with NucBlue fixed cell ReadyProbes reagent (Thermo Fisher Scientific). Coverslips were mounted with Dako fluorescence mounting medium (Agilent). The confocal imaging was performed at the MPIB Imaging Facility, on a LEICA TCS SP8 AOBS confocal laser scanning microscope (Wetzlar, Germany) equipped with a LEICA HCX PL APO 63x/NA1.4 oil immersion objective using the Leica LAS X software V3.5.7.23225. Images were analyzed with Image J 1.54p (https://imagej.net/ij/)^79^.

### Gal3 FRET assay

110,000 cells per well of the Gal3 reporter cell line RC-Gal3 were dispensed into a 12-well plate. 16-24 h later, 5 µL of the ecotropic booster receptor (Takara) was added to the wells that were going to be infected with the LVs, the plate was centrifuged at 1,200 x g for 15 min at 18 °C and placed back in the incubator. After 2 hr, the aggregate transfection was performed and LVs added to the cells. TauRD-YFPAgg cell pellets were lysed in 1% Triton X-100/PBS (Thermo Fisher Scientific) supplemented with protease inhibitor cocktail (MERCK, EDTA-free) and benzonase (prepared in-house) and kept on ice for 20 min. Protein concentration was quantified using BCA Protein Assay (Thermo Fisher Scientific). 55 ng of tau PFFs (see “Protein expression, purification and aggregation”) or 0.2 µg total protein from tau aggregate-containing cell lysate were mixed with a mixture of 50 µl Opti-MEM (Thermo Fisher Scientific) and 1.6 µL lipofectamine 3000 reagent (Thermo Fisher Scientific) and incubated for 20 min at room temperature. The mixtures were then added to the cells. 2.8×10^8^ physical particles of LVTauS or LVTauAgg quantified by HIV Type 1 p24 antigen ELISA 2.0 (ZeptoMetrix, Biozol) were added to the wells containing the ecotropic booster receptor and the plate was centrifuged at 1,200 x g for 15 min at 18 °C. After 24 hr, cells were collected using TrypL Express Enzyme (Thermo Fisher Scientific), fixed with 4% PFA in PBS for 10 min, washed with PBS and resuspended in PBS for analysis with an Attune NxT flow cytometer v.6.2.3337.1 (Thermo Fisher Scientific). To measure the mClover3 and FRET fluorescence signals, cells were excited with 488 nm laser light and fluorescence was determined using 530/30 and 590/40 filters, respectively. Data processing was performed using MatLabR2025a. The code for data processing was deposited (https://github.com/csitron/MATLAB-Programs-for-Flow-Cytometry).

### Protein expression, purification and aggregation

TauRD (tau residues 244-371, C291A/P301L/C322A/V337M) and tauRD (tau residues 244-371, P301L/V337M)-YFP were expressed and purified as described before^31^. A detailed differentiation protocol is available from protocols.io (dx.doi.org/10.17504/protocols.io.x54v9p6p1g3e/v1).

TauRD aggregation: 100 μL of 10 μM tauRD, 2.5 μM heparin (Merck, H3393), 2 mM MgCl_2_ in PBS 1x pH 7.2 were incubated at 37 °C with constant agitation (850 rpm) in a thermomixer (Eppendorf). After 1 h the sample was aliquoted, flash-frozen in liquid nitrogen, and stored at −80 °C until needed^31^.

PR(D25N) was expressed as N-terminal His6-ubiquitin fusion protein in *Escherichia coli* BL21(DE3) cells transformed with the pHUE-PR(D25N)-FLAG plasmid via IPTG induction. The cell pellet from 5 L culture was resuspended in lysis buffer (50 mM Tris-HCl pH 8.0, 150 mM NaCl) supplemented with Complete EDTAfree protease inhibitor cocktail (MERCK) and Sm DNase 50 U ml^−1^. Cells were lysed using a French press (3x cycles at 10,000 kPa) and the lysate was cleared by centrifugation (45 min, 40,000 × rpm at 4 °C). The supernatant was loaded onto a Ni-NTA column (15 mL matrix) equilibrated with lysis buffer. The column was washed with a step gradient with elution buffer (EB: 50 mM Tris-HCl pH 8.0, 150 mM NaCl, 500 mM imidazole) of 80 mL 2% EB, 80 mL 10% EB and 80 mL 25% EB. The His6-Ubiq-PR(D25N) protein was eluted with 60 mL 100% EB. The eluted fractions were collected and the salt concentration was reduced using a HiPrep 26/10 desalting column with 50 mM Tris-HCl pH 8.0, 150 mM NaCl buffer. The eluted protein was then incubated with Usp2 ubiquitin protease (prepared in-house, 1 mg Usp2 /100 mg protein), 4 mM β-Mercaptoethanol, 2.6% glycerol overnight at 4 °C to cleave the His_6_-ubiquitin tag. The cleavage mixture was applied to a Ni-NTA column equilibrated with 50 mM Tris-HCl pH 8.0, 150 mM NaCl buffer to remove His_6_-ubiquitin tag and the Usp2 ubiquitin protease. The flow-through containing PR(D25N) protein was collected and buffer was exchanged to 50 mM Tris-HCl pH 8.0, 20 mM NaCl buffer using a desalting column. Fractions containing PR(D25N) protein were loaded onto a MonoQ (10/100 GL). Protein was eluted using a gradient from 2 - 40% elution buffer (50 mM Tris-HCl pH 8.0, 1 M NaCl) over 120 mL. The eluted fractions were collected, concentrated by ultrafiltration using Vivaspin MWCO 3,000 (GE Healthcare) and loaded onto a Superdex 75 (HiLoad 16/600) column equilibrated with PBS pH 7.4. Fractions containing pure PR(D25N) protein were combined, concentrated by ultrafiltration using Vivaspin MWCO 3,000 (GE Healthcare), aliquoted and flash-frozen in liquid nitrogen for storage at −70 °C.

### Lentivirus production and Optiprep gradient

To produce LVs, 14×10^6^ cells were plated in a T175 flask. Next day cells were transfected with the corresponding viral plasmids using lipofectamine 3000 (Thermo Fisher Scientific). Specifically, the following mix was prepared: one tube with 2,250 µL Opti-MEM (Thermo Fisher Scientific) mixed with 108 µL lipofectamine 3000 reagent (Thermo Fisher Scientific) and another tube with 2,250 µL Opti-MEM (Thermo Fisher Scientific) mixed with 15 µg psPAX2 packing plasmid, 15 µg of pMD2.G E-MLV Env plasmid, 9 µg of pLenti-mRuby plasmid and 72 µL of P3000 reagent (Thermo Fisher Scientific). Both tubes were mixed and incubated for 20 min at room temperature. The culture medium was replaced with 12 mL of fresh Opti-MEM (Thermo Fisher Scientific) and the transfection mix. After 6 h, the transfection medium was removed and 30 mL of fresh medium was added to the cells. When using a different cell culture dish, all reagents were respectively scale down based on the surface area of the dish. 48h after transfection, the medium was collected, centrifuged at 300 x g 4 min and filtered through a 45 µm PE filter (MERCK) to remove remaining cells. The clarified medium (30 mL) was ultracentrifuged at 100,000 x g for 2 hr at 4 °C through a 20% sucrose/PBS cushion (6 mL). The pellet was resuspended in PBS 1 mM EDTA, aliquoted, flash frozen and stored at −70 °C.

To produce LVs in the presence of inhibitors, 1 µM Indinavir (MERCK) and 10 µM Ritonavir (MERCK) were added to the medium after exchange of the transfection medium and the same concentration was added to the storage buffer (PBS 1 mM EDTA). For seeding experiments, same concentration of both inhibitors was added to the medium.

LVs used for TIRF microscopy were produced using 10 cm^2^ dish previously coated with poly-D-lysin (Thermo Fisher Scientific). The transfection medium was removed 6 hr after transfection and exchanged for 10 mL fresh medium containing 0.1 µM of Janelia Fluor 646 Halo tag ligand (Promega). After 16 hr, the medium with Janelia Fluor 646 Halo tag ligand was collected and kept at 4 °C and 5 mL of fresh medium containing 5 µM 1,1′-dilinoleyl-3,3′-oxacarbocynanine perchlorate (DiO, Biotium) was added to the cells. After 4 hr, the DiO containing medium was removed and the Janelia Fluor 646 Halo tag ligand containing medium was added to the cells again. After further 24 hr the LVs were collected, pelleted and stored as described above.

To produce LVs for the fusion assay, 15 µg of the pN1_CypA_HiBiT plasmid was included in the transfection mix containing 15 µg psPAX2 and 15 µg pMD2.G E-MLV Env plasmids.

When performing the Optiprep gradient (Supplementary Fig. 1c), the LVs pellet was resuspended with 200-500 µl of PBS and layered on top of an Optiprep (Iodixanol, MERCK) gradient (5-20%). The Optiprep gradient was prepared the day before by layering 6 ml of 20% Optiprep/PBS beneath 6 ml of 5% Optiprep/PBS in a 13.2 mL Open-Top Thinwall Ultra-Clear Tube, 14 x 89mm (BECKAN COULTER) using a syringe and needle. The gradient was generated by using a Gradient Master (Biocomp, time: 2:05 min; angle: 81.5; speed: 12) and stored at 4 °C overnight. The gradient with the LV particles layered on top was subjected to ultracentrifugation at 250,000 x g for 1.5 hr at 4 °C (deceleration 9). After centrifugation, twelve fractions of 0.8 mL were collected using a Piston Gradient Fractionator coupled to an A254 nm spectrophotometer (Biocomp). For SDS-PAGE and immunoblotting, the fractions were precipitated by adding 1/25 volume of 2% Na-deoxycholate. After 15 min incubation on ice, 1/10 volume of 100% trichloroacetic acid was added, followed by 1 hr incubation on ice. The samples were then centrifuged at 20,000 x g for 30 min at 4 °C. The pellets were washed with 500 μL ice-cold acetone, sonicated in a Bioruptor sonication bath (Diagenode) (2 cycles of 30 s on – 30 s off) and centrifuged at 20,000 x g for 10 min at 4 °C. The pellets were air dried and resuspended with HU buffer (8 M urea, 5% SDS, 200 mM Tris-HCL pH 6.8, 1 mM EDTA, 0.01% bromophenol blue, 2% β mercaptoethanol) and incubated at 60 ⁰C shaking for 10 min.

For further LVs purification, fractions containing the p24 viral protein were combined, diluted with PBS to a final volume of 10 mL and ultracentrifuged at 100,000 x g for 2 hr at 4 °C through a 20% sucrose/PBS cushion (2 mL). Pelleted particles were resuspended in PBS 1mM EDTA, aliquoted, flash frozen and stored at −70 °C.

To produce LVs for generating stable cell lines, 4.5×10^6^ HEK293T cells for lentiviral packaging (Lenti-X 293T cell line, Takara) were plated in a 10 cm^2^ dish. Next day cells were transfected with the corresponding viral plasmids using lipofectamine 3000 (Thermo Fisher Scientific). Specifically, the following mix was prepared: one tube with 750 µL Opti-MEM (Thermo Fisher Scientific) mixed with 36 µL lipofectamine 3000 reagent (Thermo Fisher Scientific) and another tube with 750 µL Opti-MEM (Thermo Fisher Scientific) mixed with 5 µg psPAX2 packing plasmid, 5 µg of pMD2.G plasmid, 10 µg of pCW-TetOffTauRD(P301L/V337M)-Halotag plasmid, pLenti-mCat1 plasmid or pFhSynW2 TauRD (P301L/V337M)-EYFP^31^ and 24 µL of P3000 reagent (Thermo Fisher Scientific). Both tubes were mixed and incubated for 20 min at room temperature. The culture medium was replaced with 4 mL of fresh Opti-MEM (Thermo Fisher Scientific) and the transfection mix. After 6 h, the transfection medium was removed and 10 mL of fresh medium was added to the cells. 48h after transfection, the medium was collected, centrifuged at 300 x g 4 min and filtered through a 45 µm PE filter (MERCK) to remove remaining cells. LVs were precipitated from the clarified medium (10 mL) using the Lenti-X concentrator (Takara) following manufacturer’s instructions. Pellets were resuspended with 100 µL PBS, aliquoted, flash frozen and stored at −70 °C.

### Proteinase K digestion

Optiprep gradient purified LV particles were incubated with or without 0.1% Triton X-100/PBS for 20 min on ice, followed by incubation with or without proteinase K (MERCK). Approximately 13.5×10^10^ physical particles of LVs quantified using the HIV Type 1 p24 antigen ELISA 2.0 (ZeptoMetrix, Biozol) were digested in a total volume of 10 µL with 500 ng of proteinase K. The samples were digested for 30 min at 22 °C. The digestion was stopped by adding 5 µM PMSF, boiled for 5 min in NuPAGE LDS Sample Buffer (4X) buffer (Thermo Fisher Scientific) containing 100 mM DTT and proteins separated by electrophoresis.

### Gel electrophoresis and immunoblotting

For sodium dodecyl sulfate-polyacrylamide gel electrophoresis (SDS-PAGE), LVs were lysed with 1% Triton X-100/PBS, Complete EDTA-free protease inhibitor cocktail (MERK) and benzonase for 20 min on ice and boiled 5 min in NuPAGE LDS Sample Buffer (4X) buffer (Thermo Fisher Scientific) containing 100 mM DTT or directly boiled in sample buffer. To detect phosphoTau, Tris-buffered saline (TBS) was used in the lysis buffer instead of PBS and it was also complemented with PhosSTOP (MERCK).

Samples were separated by electrophoresis on NuPAGE 4–12% or 12% Bis-Tris SDS gels (Thermo Fisher Scientific) using NuPAGE MES SDS running buffer (Thermo Fisher Scientific) at 140 V. Proteins were transferred using a Power Blotter XL (Invitrogen) with Select transfer stacks nitrocellulose (Invitrogen). Membranes were blocked for at least 1 hr with TBS containing 0.05% Tween 20 (0.05% TBS-Tween) and 5% low fat milk. Immunodetection was performed using Anti-Human Tau/Repeat Domain (2B11) Mouse IgG MoAb (TECAN, JP10237, 1/100 dilution), Human Immunodeficiency Virus type 1 (HIV-1) p24 / Capsid Protein p24 Antibody (Sin Biological, 11695-R002, 1/1000 dilution), ANTI- FLAG M2 antibody, Mouse monoclonal (MERCK, F1804, 1/1000 dilution), Anti-GAPDH Antibody (MERCK, MAB374, 1/1000 dilution), Tau (phospho Ser356) antibody (GeneTex, GTX50165, 1/1000 dilution) and Anti-LgBiT Monoclonal Antibody (Promega, N7100, 1/1000 dilution). Anti-Mouse IgG (whole molecule)-Peroxidase antibody produced in goat (Merck, A4416, 1/5000 dilution), mouse anti-rabbit IgG-HRP (Santa Cruz Biotechnology, sc-2357, 1/5000 dilution) were used as secondary antibodies. Immobilon Classico Western HRP substrate (MERCK), Immobilon Forte Western HRP substrate (MERCK) or SuperSignal West Atto Ultimate Sensitivity Substrate (Thermo Fisher Scientific) were used for detection with an Amersham ImageQuant 800 GxP employing Amersham ImageQuant 800 control software 2.1.0. Full scan blots are provided in the Source Data. Quantification of band intensities was performed using Image J 1.54p (https://imagej.net/ij/)^79^.

### TIRF microscopy and analysis

LV particles were digested with proteinase K (see “Proteinase K digestion”). The samples were fixed with 2% PFA for 10 min, diluted 1/10 with ammonium chloride and transferred to a µ-Slide 8 Well high chamber (Ibidi). Total Internal Reflection Fluorescence (TIRF) microscopy was performed at the MPIB Imaging Facility, on a Thunder inverted widefield microscope equipped with an sCMOS camera Leica DFC9000 GTC using a Leica HC PL APO 63x/NA 1.47 oil immersion objective. Fluorescence channels were GFP (455 – 495 nm, Em 505-555 nm, exposure time 200 ms) for DiO signal and Cy5 (Ex 600-660 nm, Em 663-738 nm, exposure time, 500 ms) for Janelia Fluor 646 using the Leica Application Suite X 3.9.28093. Images were analyzed with Image J 1.54p^79^. Quantification of the tau signal was performed using the software ImageJ^79^, complemented with all the default plugins provided by FIJI^80^ and with the additional plugins

FeatureJ (http://imagescience.org/meijering/software/featurej/) and MorphoLibJ^81^ from the update sites ImageScience and IJPB-plugins, respectively. A script was written to repeat consistently the analysis procedure on all the stack images. The location of the viral particles was determined from the DiO signal. First, the background of the corresponding channel was removed using the rolling ball algorithm (Subtract Background, radius = 25 pixels). The particles were detected by applying a Laplacian of Gaussian filter (FeatureJ Laplacian, smoothing scale = 1 pixel) to enhance spot-like structures, followed by automated thresholding using the Triangle algorithm and conversion to a binary mask. Small spurious objects smaller than 0,05 µm^2^ were removed (MorphoLibJ, Area Opening, size = 5 pixels). Individual particles were then labeled (MorphoLibJ, Connected Component Labeling, 8-connectivity), and the labels were used to measure the average tau intensity of each viral particle from the corresponding channel (MorphoLibJ, IntensityMeasurements). The percentage of tau positive particles in each sample was computed as the percentage of particles with intensity above the background level, which was estimated using the statistics of the particles tau intensity in the LVs control sample. The threshold was set to three standard deviations above the mean. At least 12,000 particles per biological replicate were analyzed (the average number of particles analyzed per sample was around 19,000).

### Immunoprecipitation

For anti-GFP immunoprecipitation followed by in-gel mass spectrometry, LVs samples were lysed with 1% Triton X-100/PBS, 5X Complete EDTAfree protease inhibitor cocktail (MERCK), PhosSTOP (MERCK) and benzonase (prepared in-house) and incubated 20 min on ice. Total protein was quantified using the Pierce Dilution-Free Rapid Gold BCA Protein Assay (Thermo Fisher Scientific). 25 µL of ChromoTek GFP-Trap Magnetic Agarose beads (Proteintech) or ChromoTek Binding Control Magnetic Agarose beads (Proteintech) were washed with 1% Triton X-100/PBS. Beads were incubated with 75 µg of total protein overnight at 4 °C with rotation. The beads were washed 3x with PBS and eluted with 50 µL NuPAGE LDS Sample Buffer (4X) buffer (Thermo Fisher Scientific) containing 100 mM DTT. Samples were separated by electrophoresis on NuPAGE 12% Bis-Tris SDS gels (Thermo Fisher Scientific) using NuPAGE MES SDS running buffer (Thermo Fisher Scientific) at 140 V. Gel were stained with Der Blaue Jonas ultrafast protein stain (2BScientific). Gels were then excised at the Mass Spectrometry Facility (RRID:SCR_025745) at the Max Planck Institute of Biochemistry (see “Mass spectrometry”).

For anti-GFP immunoprecipitation from TauRD-YFPAgg and TauRD-YFPS cells producing LVs followed by mass spectrometry, cell pellets were lysed with 1% Triton X-100/PBS, Complete EDTAfree protease inhibitor cocktail (MERCK), PhosSTOP (MERCK) and benzonase (prepared in-house) and incubated 20 min on ice. Total protein was quantified using the Pierce Rapid Gold BCA Protein Assay (Thermo Fisher Scientific). 500 µg total protein diluted in a total volume of 500 µL lysis buffer without DNase were incubated overnight at 4 °C with rotation with 25uL of GFP-Trap Magnetic Agarose beads (Proteintech) previously washed with lysis buffer. After overnight incubation, the beads were washed 3x with PBS and prepared for mass spectrometry using the iST Sample Preparation Kit (Preomics) following manufacturer’s instructions.

For anti-GFP immunoprecipitation from TauRD-YFPAgg cells expressing PR(D25N)-FLAG or NC-FLAG or TauRD-YFPAgg cells producing LVs followed by immunoblotting, cell pellets were lysed with 1% Triton X-100/PBS, 5% glycerol, Complete EDTAfree protease inhibitor cocktail (MERCK) and benzonase (prepared in-house) and incubated 20 min on ice. Total protein was quantified using the Pierce Dilution-Free Rapid Gold BCA Protein Assay (Thermo Fisher Scientific). 12.5 µL of ChromoTek GFP-Trap Magnetic Agarose (Proteintech) were washed with binding buffer (25 mM Hepes pH 7.2, 100 mM KCl, 0.05% Triton X-100, 0.01% Tween) containing 5% glycerol and blocked with binding buffer containing 5% glycerol and 3% bovine serum albumin for 2 hr at room temperature with rotation. Beads were washed with binding buffer with 5 % glycerol and incubated with 4 mg total protein for 2 hr at room temperature with rotation. Beads were then washed with binding buffer binding buffer 5 % glycerol for 10 min with rotation, washed seven times with wash buffer (25 mM Hepes pH 7.2, 300 mM KCl, 0.1% TritonX-100, 0.02% Tween) containing 5 % glycerol for 2 min with rotation each wash, one time with wash buffer for 2 min with rotation and one time with binding buffer for 2 min with rotation. The samples were then eluted with 50 µL NuPAGE LDS Sample Buffer (4X) buffer (Thermo Fisher Scientific) containing 100 mM DTT and separated by electrophoresis. 20 µg of total protein were loaded as input. In the case of anti-GFP immunoprecipitation from TauRD-YFPAgg cells expressing PR(D25N)- FLAG or NC-FLAG 5% glycerol was omitted.

For anti-FLAG immunoprecipitation, 5 µM of recombinant PR-FLAG was mixed and incubated with 5 µM of recombinant tauRD monomers or aggregates in a total volume of 200 µL in binding buffer (25 mM Hepes pH 7.2, 100 mM KCl, 0.05% Triton X-100, 0.01% Tween) for 30 min at 25 °C. Aggregates were previously sonicated using a Bioruptor sonication bath (Diagenode) (25 cycles of 5 s on – 5 s off). 20 µL of Dynabeads Protein G for Immunoprecipitation (Thermo Fisher Scientific) were washed with antibody binding buffer from the Dynabeads Protein G Immunoprecipitation kit (Thermo Fisher Scientific). Beads were incubated with 3 µg of Anti-FLAG M2 antibody, Mouse monoclonal (Thermo Fisher Scientific, F1804) in 160 µL antibody binding buffer for 30 min at room temperature with rotation and subsequently blocked with binding buffer containing 3% bovine serum albumin for 2 hr at room temperature with rotation. Beads were washed two times with binding buffer for 3 min at room temperature with rotation and then incubated with the protein mix for 1 hr at room temperature with rotation. Beads were then washed with binding buffer for 10 min with rotation, washed eight times with wash buffer (25 mM Hepes pH 7.2, 300 mM KCl, 0.1% TritonX-100, 0.02% Tween) for 2 min with rotation each wash and one time with binding buffer for 2 min with rotation. The samples were then eluted with 45 µl NuPAGE LDS Sample Buffer (4X) buffer (Thermo Fisher Scientific) containing 100 mM DTT and separated by electrophoresis.

### Mass spectrometry

#### Sample preparation for proteomics

For mass spectrometry of anti-GFP immunoprecipitation from TauRD-YFPAgg and TauRD-YFPS cells producing LVs samples were prepared as described in “Immunoprecipitation”.

For in-gel mass spectrometry, samples were prepared as described in “Immunoprecipitation”. The gel pieces of interest (Supplementary Fig. 3a) were sliced and washed multiple times with 150 µL of destaining buffer (25 mM ammonium bicarbonate, 50% ethanol) and then dehydrated with 150 µL of 100% ethanol. After removing the ethanol, the gel pieces were dried using vacuum centrifugation. Next, 50 µL of digestion buffer (25 mM Tris-HCl, 10% acetonitrile, 10 ng/µL trypsin) was added. The mixture was incubated on ice for 20 min, followed by the addition of 50 µL of 25 mM ammonium bicarbonate buffer, and the gel pieces were incubated overnight at 37 °C. Peptides in the supernatant were collected, and additional peptides were extracted from the gel pieces through repeated incubation at 25 °C in 100 µL of extraction buffer (3% TFA, 30% acetonitrile), followed by centrifugation and collection of the supernatants. Finally, the gel pieces were dehydrated by incubating twice at 25 °C in 100 µL of 100% acetonitrile, and the supernatant was combined with the supernatants from the previous extraction steps. Acetonitrile was removed by vacuum centrifugation, and 70 µL of 2 M Tris-HCl, 10 mM tris(2-carboxyethyl)phosphine (TCEP), and 40 mM 2-chloroacetamide (CAA) were added. After a 30-minute incubation at 37 °C, the peptides were acidified to 1% TFA and purified using SDB-XC stage-tips.

For LVs proteome analysis by mass spectrometry, LVs samples were lysed with 1% Triton X-100/PBS, 5X Complete EDTAfree protease inhibitor cocktail (MERCK), PhosSTOP (MERCK) and benzonase (prepared in-house) and incubated 20 min on ice. Total protein was quantified using the Pierce Dilution-Free Rapid Gold BCA Protein Assay (Thermo Fisher Scientific). 5 µg of total protein were precipitated by adding 1/25 volume of 2% Na-deoxycholate. After 15 min incubation on ice, 1/10 volume of 100% trichloroacetic acid was added, followed by 1 hr incubation on ice. The samples were then centrifuged at 20,000 x g for 30 min at 4 °C. The pellets were washed with 500 μl ice-cold acetone, sonicated in a Bioruptor sonication bath (Diagenode) (2 cycles of 30 s on – 30 s off) and centrifuged at 20,000 x g for 10 min at 4 °C. The pellets were air dried. Cell pellets were resuspended in 400 µL of preheated SDC buffer containing 1% sodium deoxycholate (SDC; Sigma-Aldrich), 40 mM 2-chloroacetamide (CAA; Sigma-Aldrich), 10 mM tris(2-carboxyethyl)phosphine (TCEP; Thermo Fisher Scientific), and 100 mM Tris (pH 8.0). Samples were incubated at 95 °C for 5 min, followed by ultrasonication for 10 min using a Bioruptor (Diagenode). This heating and sonication cycle was repeated once. Samples were then diluted 1:1 with water and subjected to enzymatic digestion: first with 1 µg LysC for 1.5 h at 37 °C, followed by overnight digestion at 37 °C with 1 µg trypsin (Promega). The resulting peptide mixture was acidified with trifluoroacetic acid (MERCK) to a final concentration of 1%. Approximately 200 ng of peptides were loaded onto Evotips (Evotip Pure, Evosep).

#### LC-MS/MS data acquisition

For mass spectrometry of anti-GFP immunoprecipitation from TauRD-YFPAgg and TauRD-YFPS cells producing LVs and in-gel mass spectrometry, samples were analyzed on an Easy nLC-1200 nanoHPLC system (Thermo) coupled to a Q-Exactive HF Orbitrap mass spectrometer (Thermo) using Xcalibur (version 4.2.47) and Tune (version 2.11). Peptides were separated on home-made spray-columns (ID 75 μm, 30 cm long) packed with 1.9 μm C18 particles (Reprosil-Pur C18-AQ, Dr Maisch GmbH) using a stepwise 44 min (gel separated samples) or 135 min gradient between buffer A (0.1% formic acid in water/2% acetonitrile) and buffer B (0.1% formic acid in 80% acetonitrile/20% water). Samples were loaded on the column by the nanoHPLC autosampler at a pressure of 800 bar. No trap column was used. The HPLC flow rate was set to 0.25 μL per min during analysis. MS/MS analysis was performed with standard settings using cycles of 1 high resolution (120000 FWHM setting) MS scan followed by MS/MS scans (resolution 17500 FWHM setting) of the 7 most intense (gel separated samples) or 10 most intense ions with charge states of 2 or higher.

For LVs proteome analysis by mass spectrometry, peptides were eluted from Evotips onto a 15 cm PepSep C18 column (1.5 µm; Bruker Daltonics) using the Evosep One HPLC system (Evosep). The column temperature was maintained at 50 °C, and peptide separation was performed using the 30 samples-per-day (SPD) method. Eluted peptides were directly ionized and introduced into a timsTOF HT mass spectrometer (Bruker) via electrospray ionization. Data acquisition was performed in data-independent acquisition (DIA) PASEF mode via timsControl. Mass spectrometry covered a scan range of 100–1700 m/z, and ion mobility ranged from 1/K0 = 0.70 to 1.30 Vs·cm². The dual TIMS analyzer utilized equal ion accumulation and ramp times of 100 ms each, with a spectra rate of 9.52 Hz. For DIA-PASEF scans, the mass scan range was 350.2–1199.9 Da, and ion mobility ranged from 1/K0 = 0.70 to 1.30 Vs·cm². Collision energy was linearly ramped based on ion mobility, from 45 eV at 1/K0 = 1.30 Vs·cm² to 27 eV at 1/K0 = 0.85 Vs·cm². A total of 42 DIA-PASEF windows were acquired per TIMS scan, with switching precursor isolation windows, resulting in an estimated cycle time of 2.21 seconds.

#### Mass spectrometry data analysis

For mass spectrometry data analysis of the anti-GFP immunoprecipitation from TauRD-YFPAgg and TauRD-YFPS cells producing LVs, protein identification label-free quantitation was performed using MaxQuant^82^ (version 2.7.5.0) using default settings, except that label-free quantitation (LFQ) was enabled with normalization type “Classic”, “match-between-runs” was enabled and iBAQ-based quantitation was enabled. The human sequences of UNIPROT (version 2026-01-07) and a custom database of the viral proteins (including tauRD-YFP, Gag, E-MLV Env and mRuby. GagPol was omitted due to its expected low quantity and high redundancy with Gag sequence) were used as database for protein identification. MaxQuant used a decoy version of the specified UNIPROT database to adjust the false discovery rates for proteins and peptides below 1%.

For in-gel mass spectrometry, protein identification label-free quantitation was performed using MaxQuant^82^(version 2.1.4.0) using default settings, except that label-free quantitation (LFQ) was enabled with normalization type “none” and that iBAQ-based quantitation was enabled. The human sequences of UNIPROT (version 2024-01-18) and a custom database with viral proteins (including tauRD-YFP, MA/p17, CA/p24, SP1, NC/p7, SP2, p6Gag, SP2Pol, p6Pol, PR/p12, RT/p51, p15Pol, RT/p66, IN/p32, E-MLV Env Extracellular, E-MLV Env Transmembrane, E-MLV Env Cytoplasmic and mRuby) were used as database for protein identification. MaxQuant used a decoy version of the specified UNIPROT database to adjust the false discovery rates for proteins and peptides below 1%.

For LVs proteome analysis, raw data were analyzed using Spectronaut 20.2 in directDIA+ (library-free) mode, applying standard settings. The peak list was compared to a predicted library from the human UniProt database (downloaded in 2023, including both TrEMBL and SwissProt entries) and the viral proteins (including tauRD-YFP, Gag, E-MLV Env and mRuby. GagPol was omitted due to its expected low quantity and high redundancy with Gag sequence). To detect individual viral proteins the peak list was compared to a predicted library from the human UniProt database (downloaded in 2023, including both TrEMBL and SwissProt entries) and the viral proteins (including tauRD-YFP, MA/p17, CA/p24, SP1, NC/p7, SP2, p6Gag, SP2Pol, p6Pol, PR/p12, RT/p51, p15Pol, RT/p66, IN/p32, E-MLV Env Extracellular, E-MLV Env Transmembrane, E-MLV Env Cytoplasmic and mRuby). Cysteine carbamidomethylation was set as a fixed modification, while methionine oxidation and N-terminal acetylation were used as variable modifications. Protein quantification was performed across samples using label-free quantification (MaxLFQ) at the MS2 level.

Mass spectrometry raw data were submitted to the ProteomeXchange repository (See “Data availability”) and detailed analysis are included in the Source Data.

### Fusion assay

The fusion assay was performed similarly to previously described^41,42^. 20,000 HEK293T mCat1 LgBiT cells were dispensed in a well of a white 96 well plate with flat bottom (Greiner) in medium solution containing 1x Nano-Glo Endurazine™ Live Cell Substrate (Promega) and 0.1x DrkBiT Elution Peptide (Promega). The volume of LVs added to the fusion assay was calculated by normalization based on HiBiT incorporation using the Nano-Glo HiBiT Lytic Detection System (Promega) according to the manufacturer’s instructions. Briefly, different volume (1 µL, 2 µL, 4 µL and 6 µL) of LV particles included in the linear range of the detection limit were used. Luminescence was measure using a CLARIOstar plate reader (BMG Labtech, emission 470-80) with the ClarioStar software (version 5.70 R3). The volume corresponding to assay 2 × 10^6^ relative light units of each LV sample was added to the 96 well plate for the fusion assay. The plate was then centrifuged at 1,000 x g for 2 hr at 12 °C and after that immediately placed in a CLARIOstar plate reader (BMG Labtech) at 37 °C and 5% CO_2._ Luminescence (emission 470-80) was monitored every 2 min. When LVs produced in the presence of PI were used, the PI were added to the medium at the same concentration as for production and storage (see “Lentivirus production and Optiprep gradient”). HEK293T mCat1 and HEK293T LgBiT cell lines were used as controls.

### Immunostaining for flow cytometry

500,000 cells were pelleted at 1,000 x g for 4 min, washed with staining solution (1% BSA/PBS) and incubated on ice for 30 min with staining solution containing 2.5 µL of the PE anti-mouse Slc7a1 (Cat-1, ERR) antibody (Biolegend, 150504) in 100 µL staining solution. Cells were centrifuged and the pellet was washed 2 times with staining solution. The cells were resuspended in PBS and analyzed in an Attune NxT flow cytometer (Thermo Fisher Scientific). Cells were excited with 561 nm laser light and fluorescence was determined using the 585/15 filter. Data processing was performed using FlowJo V10 and V10.10.0 software (FlowJo LLC).

### Statistical analysis

Statistical analysis was performed with GraphPrism10 v.10.6.0 (Dotmatics). The sample sizes given in the figure legends describe measurements taken from distinct, biological replicates. Two-tailed Student’s t-test was used for comparing two groups. One-way analysis of variance (ANOVA) with Tukey’s or Dunnet’s post hoc test were used for multiple comparisons. Exact *P* values are provided in the Source Data.

## Acknowledgements

We thank John Briggs, Dominik Hrebík, Dorothy Y. Zhao, Daniel Bollschweiler and Tillman Schäfer for helpful discussion; Cole Sitron for critical revision of the manuscript, sharing materials and the code for Gal3 flow cytometry data processing; Zhenying Liu and Itika Saha for sharing materials; Nadine Wischnewski for protein purifications. We thank Martin Spitaler, Markus Oster, and Giovanni Cardone from the Max Planck Institute of Biochemistry (MPIB) imaging facility (RRID:SCR_025739) for support with flow cytometry, imaging and image processing and Barbara Steigenberger and her team from the MPIB Mass Spectrometry facility (RRID:SCR_025745) for in-gel processing and proteomics analysis. This research was funded in part by the Deutsche Forschungsgemeinschaft (DFG, German Research Foundation) under Germany’s Excellence Strategy within the framework of the Munich Cluster for Systems Neurology (EXC 2145 SyNergy—ID 390857198).

## Author contributions

PY designed, performed and analyzed the experiments with support from RDMP, FHS and LSRS. RK performed mass spectrometry analysis. PY and FUH designed the project and wrote the manuscript with input from the other coauthors.

## References

1 Jucker, M. & Walker, L. C. Propagation and spread of pathogenic protein assemblies in neurodegenerative diseases. Nat Neurosci 21, 1341–1349 (2018). 10.1038/s41593-018-0238-6

2 Chiti, F. & Dobson, C. M. Protein Misfolding, Amyloid Formation, and Human Disease: A Summary of Progress Over the Last Decade. Annu Rev Biochem 86, 27–68 (2017). 10.1146/annurev-biochem-061516-045115

3 Scheres, S. H. W., Ryskeldi-Falcon, B. & Goedert, M. Molecular pathology of neurodegenerative diseases by cryo-EM of amyloids. Nature 621, 701–710 (2023). 10.1038/s41586-023-06437-2

4 Taylor, A. I. P. & Radford, S. E. Amyloid fibril polymorphism: Structural mechanisms of assembly and the links to disease. Curr Opin Struct Biol 98, 103245 (2026). 10.1016/j.sbi.2026.103245

5 Peng, C., Trojanowski, J. Q. & Lee, V. M. Protein transmission in neurodegenerative disease. Nat Rev Neurol 16, 199–212 (2020). 10.1038/s41582-020-0333-7

6 Vaquer-Alicea, J. & Diamond, M. I. Propagation of Protein Aggregation in Neurodegenerative Diseases. Annu Rev Biochem 88, 785–810 (2019). 10.1146/annurev-biochem-061516-045049

7 Hipp, M. S., Kasturi, P. & Hartl, F. U. The proteostasis network and its decline in ageing. Nat Rev Mol Cell Biol 20, 421–435 (2019). 10.1038/s41580-019-0101-y

8 De Deyn, L. & Sleegers, K. The impact of rare genetic variants on Alzheimer disease. Nat Rev Neurol 21, 127–139 (2025). 10.1038/s41582-025-01062-1

9 Livingston, G. et al. Dementia prevention, intervention, and care: 2020 report of the Lancet Commission. Lancet 396, 413–446 (2020). 10.1016/S0140-6736(20)30367-6

10 Bruno, F. et al. Alzheimer’s disease as a viral disease: Revisiting the infectious hypothesis. Ageing Res Rev 91, 102068 (2023). 10.1016/j.arr.2023.102068

11 Jamieson, G. A., Maitland, N. J., Wilcock, G. K., Craske, J. & Itzhaki, R. F. Latent herpes simplex virus type 1 in normal and Alzheimer’s disease brains. J Med Virol 33, 224–227 (1991). 10.1002/jmv.1890330403

12 Itzhaki, R. F. et al. Microbes and Alzheimer’s Disease. J Alzheimers Dis 51, 979–984 (2016). 10.3233/JAD-160152

13 Levine, K. S. et al. Virus exposure and neurodegenerative disease risk across national biobanks. Neuron 111, 1086–1093 e1082 (2023). 10.1016/j.neuron.2022.12.029

14 Eyting, M. et al. A natural experiment on the effect of herpes zoster vaccination on dementia. Nature 641, 438–446 (2025). 10.1038/s41586-025-08800-x

15 Xie, M., Eyting, M., Bommer, C., Ahmed, H. & Geldsetzer, P. The effect of shingles vaccination at different stages of the dementia disease course. Cell 188, 7049–7064 e7020 (2025). 10.1016/j.cell.2025.11.007

16 Itzhaki, R. F. Overwhelming Evidence for a Major Role for Herpes Simplex Virus Type 1 (HSV1) in Alzheimer’s Disease (AD); Underwhelming Evidence against. Vaccines (Basel) 9 (2021). 10.3390/vaccines9060679

17 Itzhaki, R. F., Golde, T. E., Heneka, M. T. & Readhead, B. Do infections have a role in the pathogenesis of Alzheimer disease? Nat Rev Neurol 16, 193–197 (2020). 10.1038/s41582-020-0323-9

18 Leblanc, P. & Vorberg, I. M. Viruses in neurodegenerative diseases: More than just suspects in crimes. PLoS Pathog 18, e1010670 (2022). 10.1371/journal.ppat.1010670

19 Eimer, W. A. et al. Phosphorylated tau exhibits antimicrobial activity capable of neutralizing herpes simplex virus 1 infectivity in human neurons. Nat Neurosci 29, 604–616 (2026). 10.1038/s41593-025-02157-0

20 Hyde, V. R. et al. Anti-herpetic tau preserves neurons via the cGAS-STING-TBK1 pathway in Alzheimer’s disease. Cell Rep 44, 115109 (2025). 10.1016/j.celrep.2024.115109

21 Wainberg, M. et al. The viral hypothesis: how herpesviruses may contribute to Alzheimer’s disease. Mol Psychiatry 26, 5476–5480 (2021). 10.1038/s41380-021-01138-6

22 Fowler, S. L. et al. Tau filaments are tethered within brain extracellular vesicles in Alzheimer’s disease. Nat Neurosci 28, 40–48 (2025). 10.1038/s41593-024-01801-5

23 Protto, V. et al. HSV-1 infection induces phosphorylated tau propagation among neurons via extracellular vesicles. mBio 15, e0152224 (2024). 10.1128/mbio.01522-24

24 Liu, S. et al. Highly efficient intercellular spreading of protein misfolding mediated by viral ligand-receptor interactions. Nat Commun 12, 5739 (2021). 10.1038/s41467-021-25855-2

25 Liu, S. et al. Reactivated endogenous retroviruses promote protein aggregate spreading. Nat Commun 14, 5034 (2023). 10.1038/s41467-023-40632-z

26 Jia, X., Yin, Y., Chen, Y. & Mao, L. The Role of Viral Proteins in the Regulation of Exosomes Biogenesis. Front Cell Infect Microbiol 11, 671625 (2021). 10.3389/fcimb.2021.671625

27 Nolte-’t Hoen, E., Cremer, T., Gallo, R. C. & Margolis, L. B. Extracellular vesicles and viruses: Are they close relatives? Proc Natl Acad Sci U S A 113, 9155–9161 (2016). 10.1073/pnas.1605146113

28 Brito, A. F. & Pinney, J. W. Protein-Protein Interactions in Virus-Host Systems. Front Microbiol 8, 1557 (2017). 10.3389/fmicb.2017.01557

29 Burnie, J. & Guzzo, C. The Incorporation of Host Proteins into the External HIV-1 Envelope. Viruses 11 (2019). 10.3390/v11010085

30 Sanders, D. W. et al. Distinct tau prion strains propagate in cells and mice and define different tauopathies. Neuron 82, 1271–1288 (2014). 10.1016/j.neuron.2014.04.047

31 Yuste-Checa, P. et al. The extracellular chaperone Clusterin enhances Tau aggregate seeding in a cellular model. Nat Commun 12, 4863 (2021). 10.1038/s41467-021-25060-1

32 Cantin, R., Diou, J., Belanger, D., Tremblay, A. M. & Gilbert, C. Discrimination between exosomes and HIV-1: purification of both vesicles from cell-free supernatants. J Immunol Methods 338, 21–30 (2008). 10.1016/j.jim.2008.07.007

33 Dettenhofer, M. & Yu, X. F. Highly purified human immunodeficiency virus type 1 reveals a virtual absence of Vif in virions. J Virol 73, 1460–1467 (1999). 10.1128/JVI.73.2.1460-1467.1999

34 Selyutina, A. et al. GS-CA1 and lenacapavir stabilize the HIV-1 core and modulate the core interaction with cellular factors. iScience 25, 103593 (2022). 10.1016/j.isci.2021.103593

35 Bester, S. M. et al. Structural and mechanistic bases for a potent HIV-1 capsid inhibitor. Science 370, 360–364 (2020). 10.1126/science.abb4808

36 Sitron, C. S. et al. alpha-Synuclein aggregates inhibit ESCRT-III through sequestration and collateral degradation. Mol Cell 85, 3505–3523 e3517 (2025). 10.1016/j.molcel.2025.08.022

37 Nonaka, T., Watanabe, S. T., Iwatsubo, T. & Hasegawa, M. Seeded aggregation and toxicity of alpha-synuclein and tau: cellular models of neurodegenerative diseases. J Biol Chem 285, 34885–34898 (2010). 10.1074/jbc.M110.148460

38 Mirbaha, H. et al. Inert and seed-competent tau monomers suggest structural origins of aggregation. Elife 7 (2018). 10.7554/eLife.36584

39 Zhu, J. et al. VCP suppresses proteopathic seeding in neurons. Mol Neurodegener 17, 30 (2022). 10.1186/s13024-022-00532-0

40 Hogan, V. & Johnson, W. E. Unique Structure and Distinctive Properties of the Ancient and Ubiquitous Gamma-Type Envelope Glycoprotein. Viruses 15 (2023). 10.3390/v15020274

41 Stacey, J. C. V. et al. The conserved HIV-1 spacer peptide 2 triggers matrix lattice maturation. Nature 640, 258–264 (2025). 10.1038/s41586-025-08624-9

42 Yamamoto, M. et al. Cell-cell and virus-cell fusion assay-based analyses of alanine insertion mutants in the distal alpha9 portion of the JRFL gp41 subunit from HIV-1. J Biol Chem 294, 5677–5687 (2019). 10.1074/jbc.RA118.004579

43 Loving, R., Li, K., Wallin, M., Sjoberg, M. & Garoff, H. R-Peptide cleavage potentiates fusion-controlling isomerization of the intersubunit disulfide in Moloney murine leukemia virus Env. J Virol 82, 2594–2597 (2008). 10.1128/JVI.02039-07

44 Zhang, X. S. et al. Different Influences of Lipofection and Electrotransfection on In Vitro Gene Delivery to Primary Cultured Cortex Neurons. Pain Physician 19, 189–196 (2016).

45 Karra, D. & Dahm, R. Transfection techniques for neuronal cells. J Neurosci 30, 6171–6177 (2010). 10.1523/JNEUROSCI.0183-10.2010

46 Alabdullah, A. A. et al. Estimating transfection efficiency in differentiated and undifferentiated neural cells. BMC Res Notes 12, 225 (2019). 10.1186/s13104-019-4249-5

47 Mudher, A. et al. What is the evidence that tau pathology spreads through prion-like propagation? Acta Neuropathol Commun 5, 99 (2017). 10.1186/s40478-017-0488-7

48 Dimitrov, D. S. Virus entry: molecular mechanisms and biomedical applications. Nat Rev Microbiol 2, 109–122 (2004). 10.1038/nrmicro817

49 Chahal, G. S., Helbig, K. J., Parton, R. G. & Monson, E. A. The Biology of Endosomal Escape: Strategies for Enhanced Delivery of Therapeutics. ACS Nano 20, 1789–1813 (2026). 10.1021/acsnano.5c18112

50 Mirbaha, H. et al. Seed-competent tau monomer initiates pathology in a tauopathy mouse model. J Biol Chem 298, 102163 (2022). 10.1016/j.jbc.2022.102163

51 Sharma, A. M., Thomas, T. L., Woodard, D. R., Kashmer, O. M. & Diamond, M. I. Tau monomer encodes strains. Elife 7 (2018). 10.7554/eLife.37813

52 Murakami, T., Ablan, S., Freed, E. O. & Tanaka, Y. Regulation of human immunodeficiency virus type 1 Env-mediated membrane fusion by viral protease activity. J Virol 78, 1026–1031 (2004). 10.1128/jvi.78.2.1026-1031.2004

53 Wyma, D. J. et al. Coupling of human immunodeficiency virus type 1 fusion to virion maturation: a novel role of the gp41 cytoplasmic tail. J Virol 78, 3429–3435 (2004). 10.1128/jvi.78.7.3429-3435.2004

54 Winston, A. & Spudich, S. Cognitive disorders in people living with HIV. Lancet HIV 7, e504–e513 (2020). 10.1016/S2352-3018(20)30107-7

55 Carra, S. et al. Virus-like particles of retroviral origin in protein aggregation and neurodegenerative diseases. Mol Aspects Med 103, 101369 (2025). 10.1016/j.mam.2025.101369

56 Tyagi, M. et al. Arc mediates intercellular tau transmission via extracellular vesicles. bioRxiv (2024). 10.1101/2024.10.22.619703

57 Frost, B. & Dubnau, J. The Role of Retrotransposons and Endogenous Retroviruses in Age-Dependent Neurodegenerative Disorders. Annu Rev Neurosci 47, 123–143 (2024). 10.1146/annurev-neuro-082823-020615

58 Frisoni, G. B. et al. Alzheimer’s disease outlook: controversies and future directions. Lancet 406, 1424–1442 (2025). 10.1016/S0140-6736(25)01389-3

59 Klimmt, J. et al. A reproducible human brain tissue model to study physiological and disease-associated microglia phenotypes. bioRxiv, 2026.2001.2020.699077 (2026). 10.64898/2026.01.20.699077

60 Bowles, K. R. et al. ELAVL4, splicing, and glutamatergic dysfunction precede neuron loss in MAPT mutation cerebral organoids. Cell 184, 4547–4563 e4517 (2021). 10.1016/j.cell.2021.07.003

61 Dannert, A. et al. A human iPSC model of tauopathies engineered for 4R tau isoform expression endogenously develops late-stage neuronal tau pathology. Sci Transl Med 18, eadu9845 (2026). 10.1126/scitranslmed.adu9845

62 Parra Bravo, C., et al. Human iPSC 4R tauopathy model uncovers modifiers of tau propagation. Cell 187, 2446–2464 e2422 (2024). 10.1016/j.cell.2024.03.015

63 Samelson, A. J. et al. CRISPR screens in iPSC-derived neurons reveal principles of tau proteostasis. Cell 189, 1517–1534 e1519 (2026). 10.1016/j.cell.2025.12.038

64 Schambach, A. et al. Lentiviral vectors pseudotyped with murine ecotropic envelope: increased biosafety and convenience in preclinical research. Exp Hematol 34, 588–592 (2006). 10.1016/j.exphem.2006.02.005

65 Lovestam, S. et al. Twelve phosphomimetic mutations induce the assembly of recombinant full-length human tau into paired helical filaments. Elife 14 (2026). 10.7554/eLife.104778

66 Kasen, A. et al. Seed structure and phosphorylation in the fuzzy coat impact tau seeding competency. Nat Commun 16, 9240 (2025). 10.1038/s41467-025-64312-2

67 Sena-Esteves, M., Tebbets, J. C., Steffens, S., Crombleholme, T. & Flake, A. W. Optimized large-scale production of high titer lentivirus vector pseudotypes. J Virol Methods 122, 131–139 (2004). 10.1016/j.jviromet.2004.08.017

68 Bajar, B. T. et al. Improving brightness and photostability of green and red fluorescent proteins for live cell imaging and FRET reporting. Sci Rep 6, 20889 (2016). 10.1038/srep20889

69 Ho, S. M. et al. Evaluating Synthetic Activation and Repression of Neuropsychiatric-Related Genes in hiPSC-Derived NPCs, Neurons, and Astrocytes. Stem Cell Reports 9, 615–628 (2017). 10.1016/j.stemcr.2017.06.012

70 Mangeot, P. E. et al. Protein transfer into human cells by VSV-G-induced nanovesicles. Mol Ther 19, 1656–1666 (2011). 10.1038/mt.2011.138

71 Jang, W., Senarath, K., Feinberg, G., Lu, S. & Lambert, N. A. Visualization of endogenous G proteins on endosomes and other organelles. Elife 13 (2024). 10.7554/eLife.97033

72 Goedhart, J. et al. Structure-guided evolution of cyan fluorescent proteins towards a quantum yield of 93%. Nat Commun 3, 751 (2012). 10.1038/ncomms1738

73 Gu, X. et al. SAMTOR is an S-adenosylmethionine sensor for the mTORC1 pathway. Science 358, 813–818 (2017). 10.1126/science.aao3265

74 Furlong, R. A., Narain, Y., Rankin, J., Wyttenbach, A. & Rubinsztein, D. C. Alpha-synuclein overexpression promotes aggregation of mutant huntingtin. Biochem J 346 **Pt** **3**, 577–581 (2000).

75 Jager, S. et al. Global landscape of HIV-human protein complexes. Nature 481, 365–370 (2011). 10.1038/nature10719

76 Catanzariti, A. M., Soboleva, T. A., Jans, D. A., Board, P. G. & Baker, R. T. An efficient system for high-level expression and easy purification of authentic recombinant proteins. Protein Sci 13, 1331–1339 (2004). 10.1110/ps.04618904

77 Saha, I. et al. The AAA+ chaperone VCP disaggregates Tau fibrils and generates aggregate seeds in a cellular system. Nat Commun 14, 560 (2023). 10.1038/s41467-023-36058-2

78 Yuste-Checa, P. et al. Structural analyses define the molecular basis of clusterin chaperone function. Nat Struct Mol Biol 32, 2035–2045 (2025). 10.1038/s41594-025-01631-4

79 Schneider, C. A., Rasband, W. S. & Eliceiri, K. W. NIH Image to ImageJ: 25 years of image analysis. Nat Methods 9, 671–675 (2012). 10.1038/nmeth.2089

80 Schindelin, J., et al. Fiji: an open-source platform for biological-image analysis. Nat Methods 9, 676–682 (2012). 10.1038/nmeth.2019

81 Legland, D., Arganda-Carreras, I. & Andrey, P. MorphoLibJ: integrated library and plugins for mathematical morphology with ImageJ. Bioinformatics 32, 3532–3534 (2016). 10.1093/bioinformatics/btw413

82 Cox, J. & Mann, M. MaxQuant enables high peptide identification rates, individualized p.p.b.-range mass accuracies and proteome-wide protein quantification. Nat Biotechnol 26, 1367–1372 (2008). 10.1038/nbt.1511

