## Supplementary Information for "Virus-mediated tau aggregate seeding in a cellular model"

Supplementary information includes 4 Supplementary Figures.

### Supplementary Figures

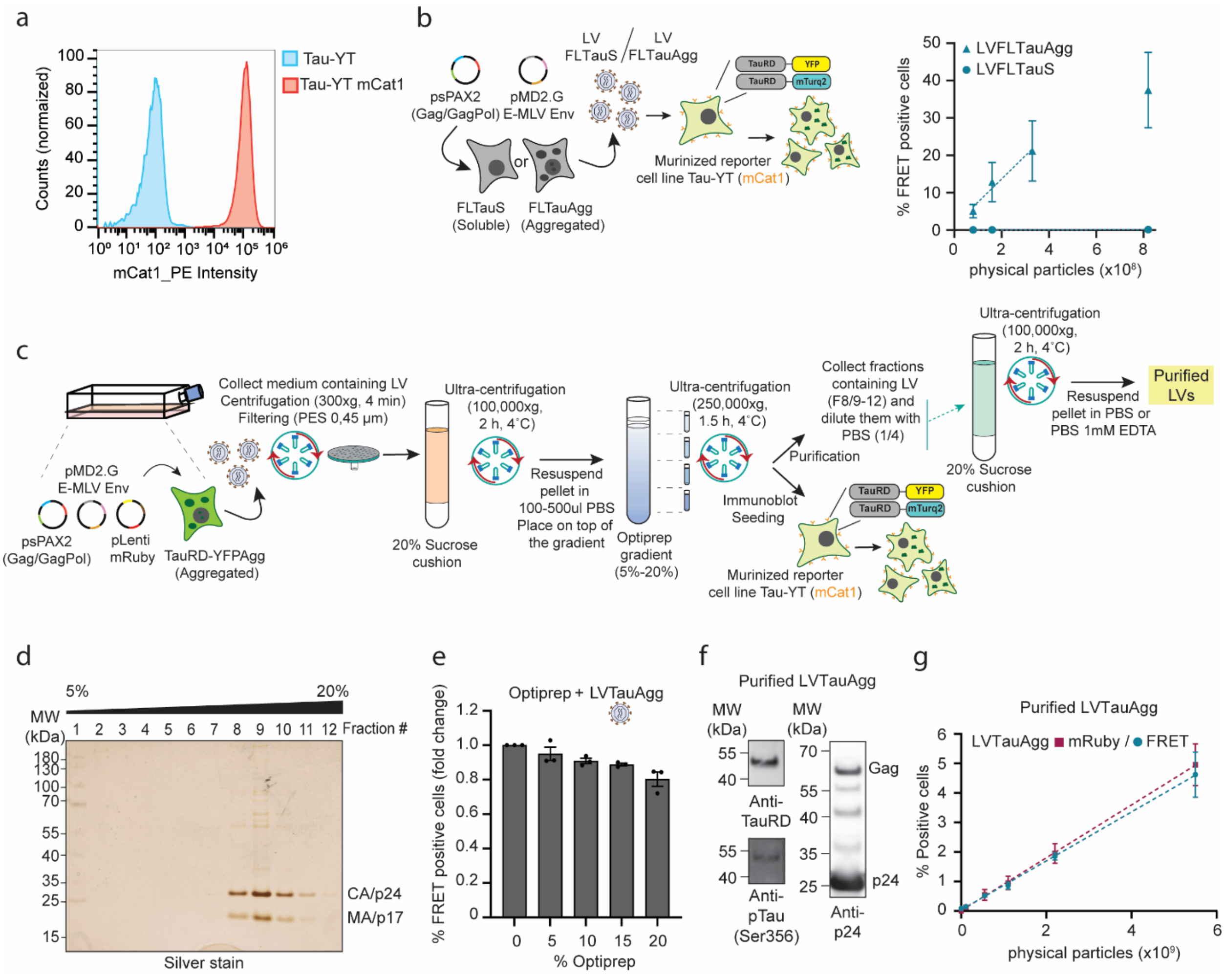


**Supplementary Figure 1. LV particle purification and seeding. a.** Quantification of mCat1 expression in tau seeding reporter cells (TauRD-YT) by immunostaining (mCat1-PE antibody) and flow cytometry. (n = 3 biological replicates). **b.** Workflow of LV production and seeding experiment. HEK293T cells overexpressing soluble (S) or aggregated (Agg) full length tau (FLTau P301L/V337M) were transfected with plasmids encoding the structural viral proteins Gag/GagPol and the ecotropic Murine Leukemia Virus envelope (E-MLV Env, murine-tropic). LVs produced in cells with soluble or aggregated FLTau (LVFLTauS and LVFLTauAgg) were added to reporter cells co-expressing TauRD fused to the FRET pair of YFP and mTurquoise2 (TauRD-YT) and murinized by expression of the murine receptor mCat1 (Tau-YTmCat1), receptor for the E-MLV Env protein (see Fig. 1a). Increasing amounts of LVFLTauS and LVFLTauAgg were added to the murinized reporter cells (Tau-YTmCat1) and induction of aggregation (FRET positive cells) was quantified by flow cytometry. Mean ± SEM (n = 3 biological replicates). **c.** Workflow of LV purification by sequential sucrose cushion and Optiprep gradient centrifugation. TauRD-YFPAgg cells were transfected with the viral plasmids encoding for the structural viral proteins Gag/GagPol, E-MLV Env protein (murine-tropic) and mRuby with the lentiviral mRNA packing signal. After 48 hours the culture medium was collected, centrifuged and filtered to remove remaining cells. The clarified medium was ultracentrifuged at 100,000 x g for 2 hours at 4 °C through a 20% sucrose cushion. The pellet was resuspended in PBS and placed on top of an Optiprep (Iodixanol) gradient (5-20%) in PBS. The gradient was centrifuged at 200,000 x g for 1.5 hours at 4 °C. The Optiprep fractions were used for immunoblotting and seeding experiments (Fig. 1d, e and Supplementary Fig. 1d, 3c-e). Fractions containing LVs were diluted in PBS and centrifuged at 100,000 x g for 2 hours at 4 °C through a 20% sucrose cushion. The pellet was resuspended in PBS 1mM EDTA, flash frozen and stored at -80 °C (Purified LVs). **d.** Representative SDS PAGE of Optiprep gradient fractions of LVTauAgg stained with silver stain. Molecular weight (MW) standards are indicated. (n = 4 biological replicates). **e.** Seeding competence of LVTauAgg mixed with PBS containing different concentrations (in % w/v) of Optiprep as control for Fig. 1e. Mean ± SEM (n = 3 biological replicates). **f.** Representative immunoblots of purified LVTauAgg. Molecular weight (MW) standards are indicated (n = 2 biological replicates). **g.** Seeding competence of purified LVTauAgg. Increasing amounts of the purified LVTauAgg were added to the murinized reporter cells (Tau-YTmCat1) and induction of aggregation (FRET positive cells, cyan) and infectivity (mRuby positive cells, maroon) was quantified by flow cytometry. Mean ± SEM (n = 3 biological replicates).

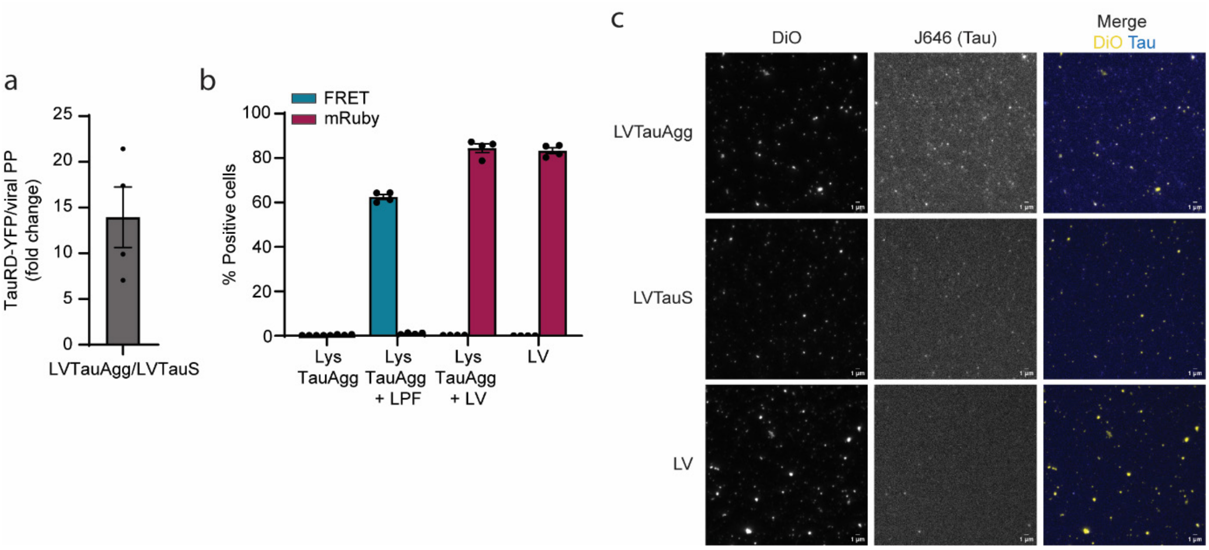


**Supplementary Figure 2. Quantification and visualization of tau in the LV particles. a.** Fold change of TauRD-YFP within the LVTauAgg particles compared to LVTauS (from Fig. 2c). Mean ± SEM (n = 4 biological replicates). **b.** Lysates from cells containing TauRD aggregates were transfected with lipofectamine (LPF) or directly added to the murinized seeding reporter cells (Tau-YTmCat1). LVs produced in naïve HEK293T cells (LV) were incubated with or without lysates from cells containing TauRD aggregates for 2 hours at 37 °C and then added to Tau-YTmCat1 cells. Induction of aggregation (FRET positive cells, cyan) and infectivity (mRuby positive cells, maroon) was quantified by flow cytometry. Mean ± SEM (n = 4 biological replicates). **c.** Representative TRIF microscopy pictures of LVTauAgg, LVTauS and control LV (which do not contain tau but are labelled with Janelia Fluor 646 and with the lipid green dye DiO) are shown (Fig. 2e-g). Scale bars, 1μm. (n = 4 biological replicates).

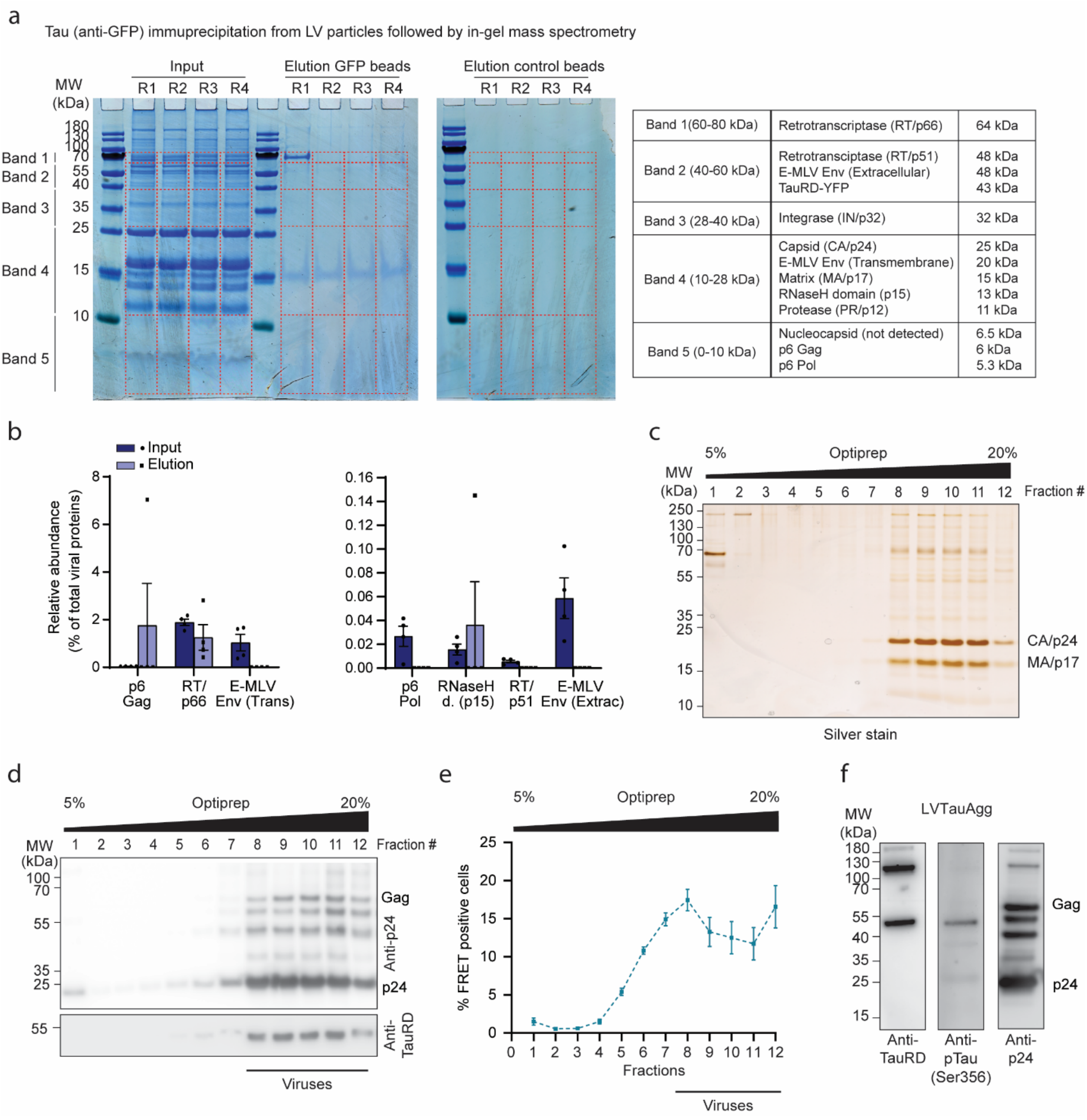


| | | |
|---|---|---|
| Band 1(60-80 kDa) | Retrotranscriptase (RT/p66) | 64 kDa |
| Band 2 (40-60 kDa) | Retrotransciptase (RT/p51)<br>E-MLV Env (Extracellular)<br>TauRD-YFP | 48 kDa<br>48 kDa<br>43 kDa |
| Band 3 (28-40 kDa) | Integrase (IN/p32) | 32 kDa |
| Band 4 (10-28 kDa) | Capsid (CA/p24)<br>E-MLV Env (Transmembrane)<br>Matrix (MA/p17)<br>RNaseH domain (p15)<br>Protease (PR/p12) | 25 kDa<br>20 kDa<br>15 kDa<br>13 kDa<br>11 kDa |
| Band 5 (0-10 kDa) | Nucleocapsid (not detected)<br>p6 Gag<br>p6 Pol | 6.5 kDa<br>6 kDa<br>5.3 kDa |

**Supplementary Figure 3. In-gel mass spectrometry and Optiprep gradient of LVTauAgg without virus-tagged mRNA. a.** Coomassie blue staining of input and elution fractions from 4 replicates (R1-4) of anti-GFP and control bead immunoprecipitations. The band excision pattern used for downstream in-gel mass spectrometry analysis is indicated. The table on the right summarizes the proteins expected to be detected in each excised band, including tau and individual viral proteins, together with their corresponding predicted molecular weights. **b.** Relative abundance expressed as percentage of total viral proteins in input and elution fractions of the anti-GFP immunoprecipitation in (a) by mass spectrometry (calculated from iBAQ intensities). The viral retrotranscriptases (RT/p66, RNAseH domain/p15, RT/p51), p6Gag, p6Pol and the extracellular and transmembrane subunits of the E-MLV Env protein, are represented in the bar graphs. Mean ± SEM (n = 4 biological replicates). **c.** Optiprep gradient of sucrose cushion-pelleted LVTauAgg without virus-tagged mRNA (detailed protocol in Supplementary Fig. 1c). Representative SDS PAGE of Optiprep gradient fractions stained with silver stain. Molecular weight (MW) standards are indicated. (n = 3 biological replicates). **d.** Representative immunoblots of the Optiprep fractions as in (c) are shown. Molecular weight (MW) standards are indicated. (n = 3 biological replicates). **e.** Seeding competence (FRET positive cells) of the different Optiprep fractions from (c) and (d). Mean ± SEM (n = 4 biological replicates). **f.** Representative immunoblots of purified LVTauAgg without virus-tagged mRNA. Molecular weight (MW) standards are indicated. (n = 2 biological replicates).

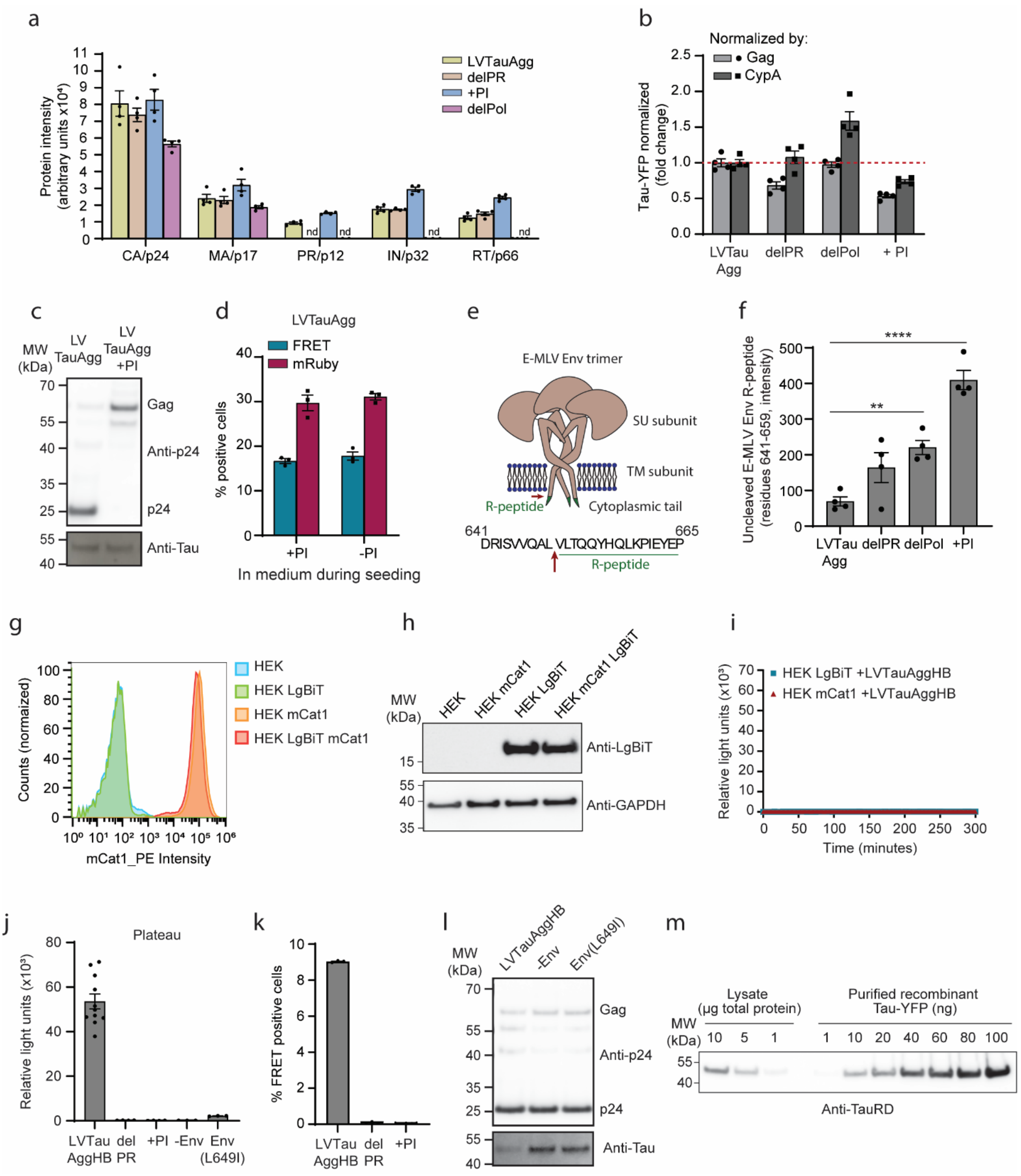


**Supplementary Figure 4. Dependency of LV fusion with recipient cells and subsequent aggregate seeding on viral protease activity. a.** Protein intensity (PG.Quantity) of the viral proteins in LVTauAgg (yellow), delPR (LVTauAgg without viral PR, orange), +PI (LVTauAgg produced in the presence of protease inhibitors, blue) and delPol (LVTauAgg without the whole Pol sequence, purple) particles detected by mass spectrometry. Mean ± SEM (n = 4 biological replicates). nd, not detected. **b.** Tau-YFP in the LVTauAgg, delPR, delPol and +PI particles quantified by mass spectrometry (PG.Quantity) and normalized by the amount of Gag polypeptide (light gray) or cyclophilin A (CypA, dark gray). LVTauAgg WT is used as reference (1). Mean ± SEM (n = 4 biological replicates). **c.** Representative immunoblots of LVTauAgg and +PI particles. Molecular weight (MW) standards are indicated. (n = 3 biological replicates). **d.** LVTauAgg particles were added to the murinized tau seeding reporter cell line (Tau-YTmCat1) in the presence or absence of PI in the culture medium. Tau seeding (FRET positive cells, cyan) and infectivity (mRuby positive cells, maroon) was quantified by flow cytometry. Mean ± SEM (n = 3 biological replicates). **e.** Simplified scheme of the ecotropic Murine Leukemia Virus envelope protein (E-MLV Env). The surface (SU) subunit, transmembrane (TM) subunit, cytoplasmic tail and cleavable R-peptide (green) are indicated. The amino acid sequence corresponds to the uncleaved peptide containing the R-peptide cleavage site detected by mass spectrometry (f). Amino acid numbers of N- and C-termini are indicated. The red arrow marks the R-peptide cleavage site. **f.** Uncleaved E-MLV Env R-peptide (e) intensity (EG.TotalQuantity) detected in LVTauAgg, delPR, delPol and +PI particles by mass spectrometry (residues 641-659). Mean ± SEM. ****$P < 0.0001$, **$P < 0.01$ according to one-way ANOVA with Dunnet's post hoc test. (n = 4 biological replicates). **g.** Quantification of mCat1 expression in the HEK293T fusion reporter cell line (HEK LgBiT mCat1) and control cell lines (HEK, HEK LgBiT and HEK mCat1) quantified by immunostaining (mCat1-PE antibody) and flow cytometry (n = 3 biological replicates). **h.** Expression of LgBiT in the HEK293T fusion reporter cell line (HEK LgBiT mCat1) and control cell lines (HEK, HEK LgBiT and HEK mCat1) detected by immunoblotting. A representative immunoblot is shown. Molecular weight (MW) standards are indicated. (n = 3 biological replicates). **i.** LVTauAggHB particles were added to control reporter cells HEK mCat1 and HEK LgBiT, and luminescence was monitored over time. Mean ± SEM (n = 6 biological replicates). **j.** Luminescence plateau reached by LVTauAggHB, delPR, +PI, -Env and Env(L649I) after 300 min of fusion assay from Fig. 4f. Mean ± SEM (n=3 biological replicates, except for LVTauAggHB n = 11; delPR and +PI n= 4). **k.** Seeding competence of LVTauAggHB, delPR (LVTauAggHB without viral PR) and +PI (LVTauAggHB produced in the presence of PI) particles. Equivalent amounts of the LVTauAgg, delPR or +PI particles normalized based on HiBiT incorporation were added to the murinized tau seeding reporter cells (Tau-YTmCat1). Induction of tau aggregation (FRET positive cells) was quantified by flow cytometry. Mean ± SEM (n = 3 biological replicates). **l.** Representative immunoblots of LVTauAggHB, -Env and Env(L649I) particles. Molecular weight (MW) standards are indicated. (n = 3 biological replicates). **m.** Representative immunoblot of total lysate of TauRD-YFPAgg cells (Fig. 1a), with purified recombinant

TauRD-YFP used as standard. Molecular weight (MW) standards are indicated. (n = 3 biological replicates).
